# Regulatory variants drive Hsp90’s effect on adaptation

**DOI:** 10.1101/2023.10.30.564848

**Authors:** Christopher M. Jakobson, José Aguilar-Rodríguez, Daniel F. Jarosz

## Abstract

The essential, stress-responsive protein chaperone Hsp90 impacts adaptation from microbes to humans. Yet the molecular mechanisms involved remain unclear and are often presumed to be driven by variation in opening reading frames. Here, we identify over 1,000 natural genotype-to-phenotype associations governed by Hsp90 at single-nucleotide resolution in *Saccharomyces cerevisiae*. Strikingly, *cis*-regulatory variants contributed to the chaperone’s effect on heredity more strongly than coding variation. Most of these mutations nonetheless impacted clients of Hsp90 or targets of its direct binding partners. The chaperone’s influence was especially potent on evolutionarily young genes, highlighting its influence on variation central to novelty. Synthetic reconstructions and genome editing revealed that Hsp90 may regulate the relationship between activity and phenotype for many dosage-sensitive genes. The unexpected role of the chaperone in buffering the effects of regulatory variants and releasing their consequences under stress suggests a mechanism for its widespread effects on evolution and development.

## INTRODUCTION

The phenotypes expressed from the genome are constantly remodeled throughout development and under stress^1^, with profound impacts on disease and evolution^2^. The stress-regulated Heat Shock Protein 90 kDa (Hsp90) chaperone has attracted keen interest as a molecular mediator of these effects^3,4^. Hsp90 is a highly expressed^5^ ‘hub of hubs’ that interacts genetically and physically^6^ with hundreds of client^7^ kinases, ubiquitin ligases, and other signaling factors^8^ that are likewise highly connected nodes of regulatory networks. Thus, Hsp90 couples the turnover, steady-state levels, and localization of much of the regulatory proteome^9,10^ to environmental stress^11^.

As a result, this chaperone has profound impacts on the phenotypic manifestation of genetic variation^11–18^. Hsp90 activity is tightly linked to oncogenesis^19,20^ and the Hsp90 chaperone network plays a central role in neurodegeneration^21^. Such effects are not limited to disease: modest impairment of Hsp90 function during development in *Drosophila melanogaster* results in pronounced changes to wings, eyes, and legs^16^, phenotypes classically associated with regulatory variation. The lineage specificity of these effects^16,22^, as well as those observed in other organisms^17^ from yeast to plants, indicates that Hsp90, in addition to its effects on disease variants, controls the fitness consequences of a substantial fraction of natural polymorphisms. Moreover, this influence is most pronounced under stress when protein quality control is compromised^23^—and when the value of phenotypic novelty is greatest.

However, owing to the limitation of existing genetic mapping technologies, the precise underlying mutations whose effects are modified by Hsp90 have remained obscure in all but a handful of instances^14^. Because Hsp90 is a chaperone of protein folding, prior studies examining the interactions between Hsp90 and, variously, gene deletions^24,25^, naturally segregating haplotypes^26^, Fanconi anemia mutations^14^, variants underlying Mendelian disorders^27^, and genotypes harboring small numbers of nearly random mutations^13^ made the pragmatic assumption that chaperone-dependent phenotypes arose primarily from coding variation. Yet the morphological and other developmental traits impacted when the chaperone’s activity is compromised are most frequently caused by *cis*-regulatory variation^28^.

Here, we take advantage of the deep conservation of the molecular activity and clientele of Hsp90^29^ and the power of quantitative genetics in the budding yeast *Saccharomyces cerevisiae* to identify, at unprecedented scale, hundreds of natural polymorphisms whose effects depend strongly on Hsp90 function. The advent of genetic mapping approaches in *S. cerevisiae* that dissect complex traits to their underlying causal variants^30,31^, in conjunction with collections of wild isolates^32,33^, genome engineering tools^34,35^, and systematic knowledge of genetic and physical interactions^36,37^ and the targets of key Hsp90 clients^38–40^, make this an ideal venue to understand the mechanisms by which the chaperone reshapes phenotypes. In contrast to prevailing assumptions, we found that even though Hsp90 is a protein chaperone, its effects were most pronounced for non-coding variation. This surprising role in the phenotypic expression of *cis*-regulatory variation provides a mechanistic explanation for Hsp90’s potent impact on the evolution of developmental phenotypes. Indeed, buffering the consequences of excursions in activity and releasing their fitness effects under stress may facilitate the accumulation of environmentally specific adaptive variation across Eukarya.

## RESULTS

### Comprehensive mapping of Hsp90-dependent phenotypes

To enable genetic mapping with high resolution and sensitivity, we employed ∼15,000 fully genotyped progeny^30^ from an F_6_ intercross between two wild *Saccharomyces cerevisiae* parents^31,41^: RM11, isolated from a California vineyard^42^, and YJM975, isolated from an immunocompromised patient^43^. Although most genes are polymorphic in this mapping panel, those encoding Hsp90—*HSC82* and *HSP82*—are not (mutations in these genes could confound mapping). To assess Hsp90’s impact on phenotypes driven by the diverse natural variants present in this cross [**Fig. 1A**], we used a potent and specific natural product inhibitor, radicicol^44^, at a dose that reduced but did not eliminate chaperone activity^18^ [**Fig. S1A**], did not perturb growth in the absence of stress [**Fig. S1B**], and did not induce a bounce-back heat shock response^26^ [**Fig. S1CD**].

**Figure 1.**
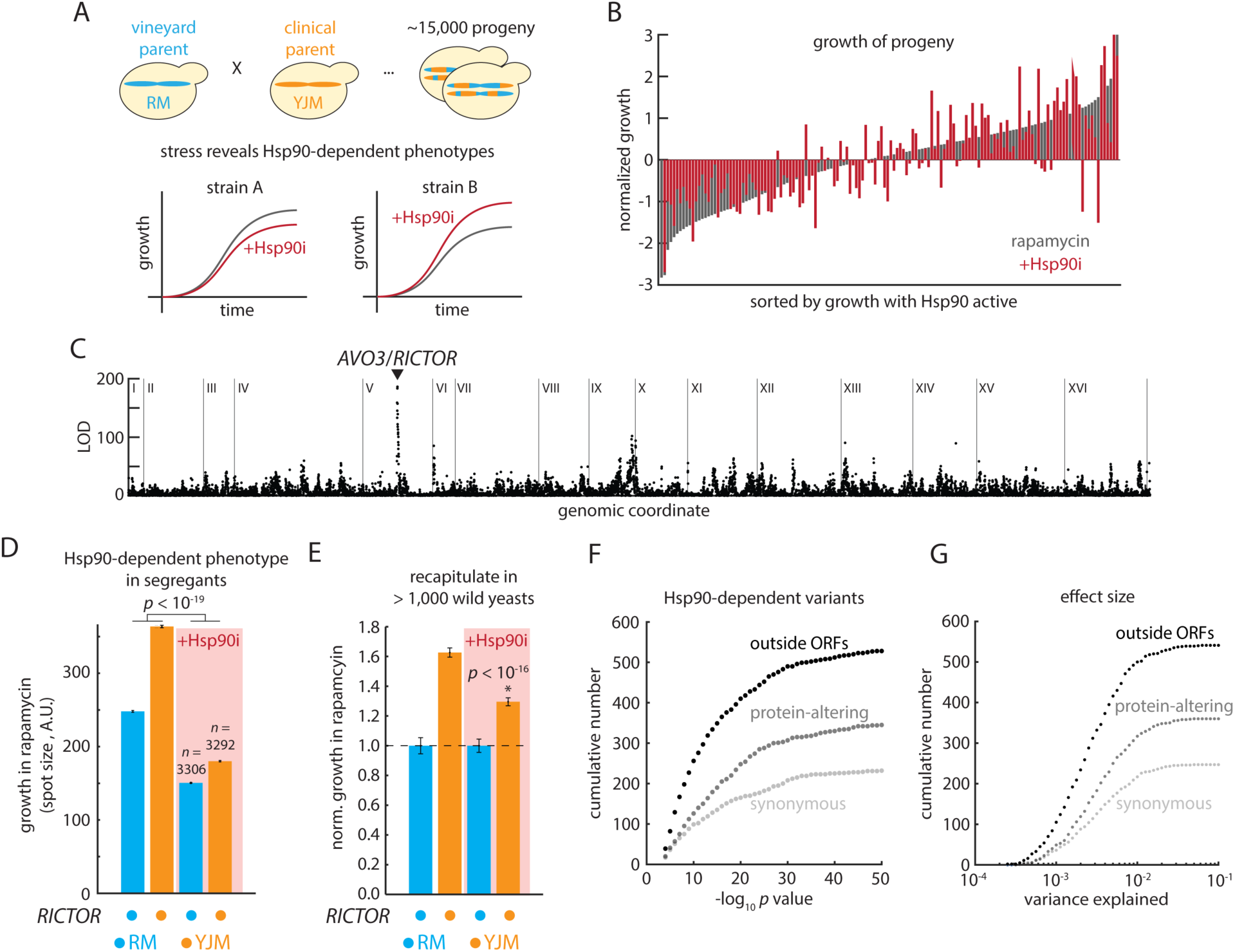
Mapping the basis of Hsp90-dependent phenotypes. (A) Schematic of genetic mapping population and hypothetical Hsp90-dependent phenotypes of different strains. (B) Rank-ordered normalized growth of 146 randomly selected progeny in rapamycin, with (red) and without (gray) radicicol. Data are normalized to mean = 0 and standard deviation = 1 amongst the F_6_ progeny in that treatment condition. (C) Manhattan plot of univariate genetic mapping of Hsp90-dependent rapamycin resistance. Loci are in genomic order, with chromosomes as indicated. LOD, log of the odds distribution. 5% FDR threshold < 5 by permutation. (D) Growth in rapamycin with (red shading) and without (no shading) radicicol of progeny bearing the sensitive (blue) and resistant (orange) alleles of Avo3/RICTOR. Data shown are mean and s.e.m. for the indicated number of progeny with each homozygous genotype; *F* test *p-*value from multivariate regression. (E) Replication of the Hsp90-dependent phenotype in wild isolates sorted by their Avo3 genotype as indicated: Avo3^Thr1342^ in blue (*n* = 152) and Avo3^Lys1342^ in orange (*n* = 806). Growth was measured with Hsp90 active (no shading) and inhibited (red shading) Data shown are mean and s.e.m. normalized to the growth of the RM11 Avo3^Thr1342^ allele; *p* value by *t* test relative to growth without radicicol. (F) Cumulative frequency plots of multivariate regression *p* values for protein-altering, synonymous, and regulatory Hsp90-modified variants. (G) Cumulative frequency plots of multivariate regression variance explained for protein-altering, synonymous, and regulatory Hsp90-modified variants. See also [**Fig. S1**].

We first examined the growth of the 15,000 diploid progeny in response to diverse environmental perturbations (including alternate carbon sources, toxic metal ions, and DNA-damaging agents, as well as antifungal drugs used clinically and in agriculture) [**Supplemental Table S1**]. Parallel experiments in the presence of radicicol revealed that Hsp90 inhibition dramatically altered the growth of many progeny [**Fig. 1B**], but did so in a manner that was highly specific to genotype and environment [**Fig. S1E**]. Coarse genetic mapping readily identified loci with Hsp90-dependent phenotypic effects logically connected to the traits we examined [**Fig. 1C**], including the rapamycin-resistance phenotype of a locus centered about the *AVO3* (*TSC11*/RICTOR^45^) gene [**Fig. 1D**] – the Avo3 protein is a key component of the TORC2 complex, a direct target of rapamycin.

To investigate whether our findings from this defined mapping panel generalized across distinct genetic backgrounds, we assessed how Hsp90 inhibition influenced growth in rapamycin across > 1,000 wild yeast isolates^33^ from diverse ecological niches and geographic origins. Resistance associated with the clinical allele of Avo3, and its Hsp90-dependence, was apparent across the >100 wild isolates that harbored this allele amidst additional variation at more than one million other loci [**Fig. 1E**]. Although genetic background can exert a strong influence on phenotype, these observations suggest that inferences about Hsp90’s relationship to the effects of specific genetic variants from our experiments may hold across a very wide range of wild isolates.

### Hsp90 is a potent buffer of regulatory variation

Integrating over 1,000,000 growth measurements and employing a multivariate mapping procedure^30^, we identified more than 5,000 Hsp90-dependent loci throughout the genome, exceeding the number previously identified by orders of magnitude [**Supplemental Table S2**], and emphasizing Hsp90’s widespread influence on the heredity of diverse biological traits. Our mapping approach provided a comprehensive picture of Hsp90-dependent heredity, revealing that up to 76.2% of Hsp90’s effect on fitness was due to changes in the phenotypic effects of genetic variation [**Fig. S1FG**]. Consistent with the minimal effect of radicicol on growth in the absence of stress, no more than 20% of the Hsp90-dependent QTLs in any environment overlapped with those identified in the no-stress condition, and even less so for the strongest-effect loci. The expression of Hsp90 itself also made a minor contribution: the lone regulatory variant near either the *HSC82* or *HSP82* genes explained only 0.08% of Hsp90-dependent growth. Moreover, only two of the Hsp90-dependent loci we identified overlapped with Hsf1 binding sites [**Supplemental Table S2**], further confirming that the heat shock response was not engaged.

We classified Hsp90-dependent loci into two broad categories. In the first, known as buffering interactions, the phenotypic effect of an allele is masked by Hsp90 and thus *increased* upon radicicol treatment, a release of cryptic variation sometimes termed decanalization^46,47^. In the second, known as potentiation interactions, the phenotypic effect of a variant is dependent on Hsp90 and thus *reduced* upon radicicol treatment (as with the transforming kinase v-Src^18^) [**Fig. S1H**]. Across the 1,154 interactions (that is, Hsp90-dependent effects on phenotype) that we identified, both behaviors were common. Buffering interactions were somewhat stronger than potentiations (median 15%; *p* < 10^-4^) [**Fig. S1I**], and we noted that Hsp90 inverted the fitness effects of 309 variants. Although they were a minority of the interactions we observed, the average effect size of such inversions was large [**Fig. S1J**] and may explain the ‘line-crossing’ Hsp90 dependence that has been reported for randomly accumulated mutations^13^. Overall, the magnitude of Hsp90’s impact on individual variants (median variance explained ∼ 0.21%) was comparable to the phenotypic contributions of the variants themselves (∼ 0.26%) [**Fig. S1K**] and, when Hsp90 was inhibited, buffered variants exhibited the largest effects [**Fig S1L**]. The majority of these effects far exceed the threshold for effective neutrality in this organism (*s* ∼ 10^-8^ - 10^-6^)^48^, establishing that Hsp90 is poised to impact the evolutionary fate of a broad swath of natural genetic variation. Consistent with reduced selection on variants that are typically buffered, allowing them to reach fixation more often, alleles buffered by Hsp90 rose to higher frequencies in *S. cerevisiae* than variants that were causal with Hsp90 active (*p* < 0.01); the same was not true of potentiated alleles (*p* = 0.27) [**Fig. S1M**].

To pinpoint the polymorphisms responsible for Hsp90-dependent traits, we performed high-resolution genetic mapping^31,41,49^ of the change in growth upon chaperone inhibition across each environmental perturbation. These nucleotide-resolution data revealed that nearly half (542 of 1,154) of the fine-mapped variants were outside of annotated open reading frames. Moreover, regulatory variants accounted for more Hsp90-dependent heredity than protein-altering variants [**Fig. 1FG; Fig. S1N**], even though only 30% of the yeast genome is noncoding and only 44% of the SNPs in our mapping panel lie outside open reading frames.

### Reconstruction by gene editing to confirm the nucleotide-resolution map

To confirm the findings from our mapping analyses, we first used genome editing to reconstruct a SNP in Avo3^50^ linked to Hsp90-dependent rapamycin resistance [**Fig. 2A**]. Statistical fine mapping suggested that a missense variant at position 1342 of the Avo3 protein was likely responsible, as indicated by a pronounced peak in the QTN score (although, notably, not in the traditional LOD metric). Examining the predicted structure^51^ of this Hsp90 client^36^, we noted a salt bridge formed by the polymorphic Lys1342 residue that appeared to stabilize the region of Avo3/RICTOR that renders TORC2 insensitive to rapamycin^52^ [**Fig. 2B**]. Consistent with this hypothesis, introduction of the alternate Thr1342 allele found in the RM11 vineyard parent by gene editing conferred pronounced rapamycin sensitivity that was attenuated by radicicol treatment [**Fig. 2C**]. By contrast, introduction of the Lys1342 allele found in the clinical parent conferred potentiated resistance [**Fig. 2C**]. We obtained similar results with radicicol and the structurally and mechanistically distinct Hsp90 inhibitor geldanamycin^53^, and neither variant strongly influenced growth in the absence of rapamycin [**Fig. S2A**], confirming that the phenotypes of these polymorphisms were transformed by Hsp90 activity.

**Figure 2.**
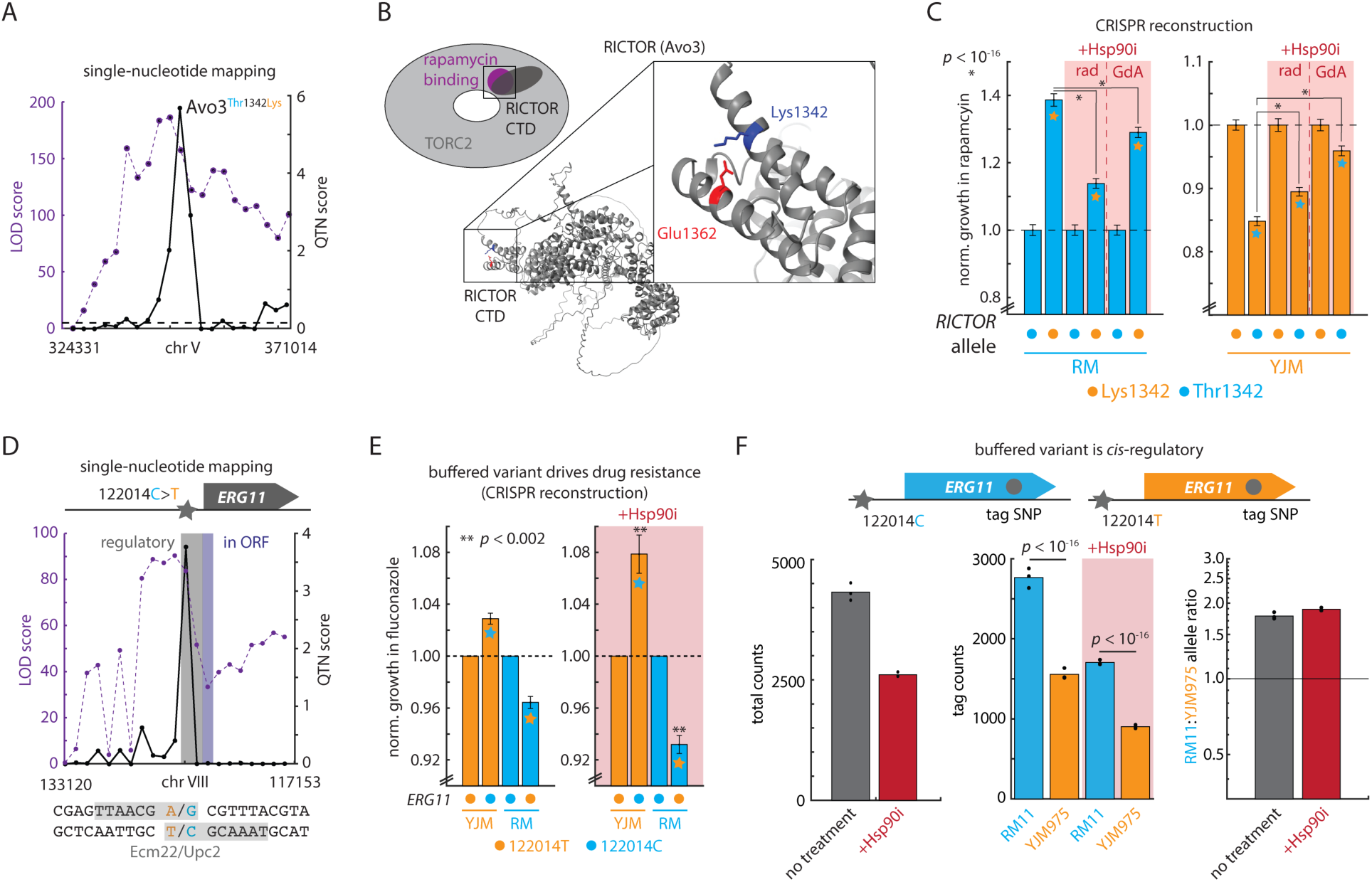
Identifying individual genetic variants underlying Hsp90-dependent phenotypes. (A) Univariate genetic mapping of Hsp90-dependent rapamycin resistance (LOD score; left ordinate) overlaid with multivariate fine-mapping (ANOVA QTN score; right ordinate) of the same phenotype, highlighting the causal variant Avo3^Thr1342Lys^. (B) Schematic of the TORC2 complex, rapamycin binding pocket, and RICTOR C-terminal domain; Alphafold2 predicted structure of Avo3, highlighting the causal residue Avo3^Lys1342^ and its proximity to Avo3^Glu1362^. (C) Growth in rapamycin of genome-edited strains with Hsp90 active (no shading) and inhibited (red shading). The right panel shows wild type RM11 Avo3^Thr1342^ strains and edited RM11 Avo3^Lys1342^ strains (orange stars); the left panel shows wild type YJM975 Avo3^Lys1342^ and edited YJM975 Avo3^Thr1342^ strains (blue stars). Hsp90 was inhibited with both radicicol (rad) and geldanamycin A (GdA). Data shown are mean and s.e.m. normalized to the wild type; *p* values by *t* test; *n* = 96 per genotype. (D) Univariate genetic mapping of Hsp90-dependent fluconazole resistance (LOD score; left ordinate) overlaid with multivariate fine-mapping (ANOVA QTN score; right ordinate) of the same phenotype, highlighting the causal variant *ERG11*^122014C>T^. Indicated below are overlapping near-cognate binding sites for Ecm22/Upc2 centered about the polymorphism. (E) Growth in fluconazole of genome-edited strains in fluconazole with (red shading) and without (no shading) Hsp90 inhibition with radicicol (+Hsp90i). Each panel shows wild type YJM975 *ERG11*^122014T^, edited YJM975 *ERG11*^122014C^ (blue stars), wild type RM11 *ERG11*^122014C^, and edited RM11 *ERG11*^122014T^ (orange stars) strains. Mean and s.e.m. are shown, normalized to the wild type; *p* values by *t* test against the same mutant strain without radicicol treatment; *n* = 384 per genotype. (F) Top: Schematic of *ERG11* alleles in the hybrid diploid; indicated are the Hsp90-dependent 122014C>T regulatory variant and the Erg11^Lys433Asn^ tag SNP. Below left: Total counts at the Erg11^Lys433Asn^ tag SNP with Hsp90 inhibited with radicicol (red) or Hsp90 active (grey). Below middle: estimated counts at Erg11^Lys433Asn^ tag SNP for the RM11 allele (blue) and the YJM975 allele (orange) with Hsp90 inhibited with radicicol (red shading) or Hsp90 active. Bars show mean and dots show replicates. Below right: Mean allelic ratio (RM11 counts/YM975 counts) at the Erg11^Lys433Asn^ tag SNP with Hsp90 inhibited with radicicol (red) or Hsp90 active (grey). Bars show mean and dots show replicates; *p* values by binomial test; *N* = 3. See also [**Fig. S2**].

Next, to better understand the surprising enrichment for Hsp90-dependent regulatory variants, we investigated a T>C variant 206 base pairs upstream of the *ERG11* transcription start site^54^ [**Fig. 2D**] linked to Hsp90-buffered resistance to the antifungal drugs tebuconazole and fluconazole, which inhibit Erg11 and are widely used in agriculture^55^ and the clinic^56^ [**Fig. S2B**]. Genomic reconstruction of the regulatory SNP recapitulated its chaperone-dependent effect on sensitivity to these mainline antifungal drugs [**Fig. 2E; Fig. S2C**]. The mutation had a negligible effect on growth in the absence of drug, regardless of Hsp90 activity [**Fig. S2D**]. The results of these genome editing studies establish that drug resistance was due to a buffering interaction between this regulatory variant and Hsp90, a chaperone of protein folding.

### Hsp90 controls the phenotypic effects of *cis-*regulatory variation

We initially hypothesized that Hsp90 might alter how these SNPs impact mRNA levels, and therefore investigated allele-specific mRNA expression in the heterozygous diploid parent of our genetic mapping panel. This approach capitalizes on polymorphisms in transcribed regions that ‘tag’ transcripts as originating from either the RM11 or YJM975 parental chromosomes [**Fig. S3A**]. A predominance of one haplotype over the other indicates the presence of a *cis*-acting regulatory variant at that gene^57^. Roughly a quarter of all non-coding variants with phenotypes that were impacted by Hsp90 (107 of 451) exhibited significant allele-specific expression (ASE) when the chaperone was inhibited (amongst ORFs with tag SNPs; Bonferroni-corrected *q <* 0.05; see **Methods**, **Fig. S3B**, and **Supplemental Table S3**). The buffered variant at *ERG11*, for instance, showed pronounced RM11-biased expression (RM:YJM allelic ratio = 1.78 ± 0.06) [**Fig. 2F**]. This *cis*-regulatory effect was consistent with the variant’s proximity to two near-cognate Ecm22/Upc2 binding motifs near the start of *ERG11* transcription [**Fig. 2D**]^58^ and our observation that the RM11 allele conferred azole resistance [**Fig. 2A**]. Hsp90 inhibition markedly decreased the overall level of *ERG11* expression, possibly because the Upc2 transcription factor associates with Hsp90^59^. Yet the biased allelic expression was preserved (1.89 ± 0.03) [**Fig. 2F**].

Indeed, Hsp90 inhibition rarely had a strong effect on ASE—only 18% of Hsp90-dependent SNPs changed in allelic ratio by more than 50%—despite often affecting expression in *trans* [**Fig. 3A**; **Fig. S3C**]. As with the *ERG11* mutation, the largest and most frequent effects we identified across the genome arose from variants upstream of the TSS, consistent with altered transcription factor recruitment or transcription initiation [**Fig. 3B**]. 123 Hsp90-dependent variants, however, were in the predicted 3’ UTR of mRNAs and 55 in the 5’ UTR [**Fig. 3C**], pointing to a significant role for post-transcriptional regulation. For example, a polymorphism that caused Hsp90-dependent resistance to both lithium and nickel stress localized to the highly structured 3’ UTR^60^ of *HAC1*, which regulates pre-mRNA splicing and levels *via* Ire1^61^. Likewise, numerous Hsp90-dependent phenotypes arose from the 3’ UTR of the cysteine desulfurase gene *NFS1*, which exhibits nutrient-dependent interactions with RNA-binding proteins^62^. Introducing a G>A polymorphism 61 base pairs downstream of *NFS1* by gene editing was sufficient to confer pronounced buffered sensitivity to fluconazole and concomitant buffered resistance to the DNA-damaging agent doxorubicin, confirming the role of this 3’ UTR in chaperone-dependent heredity [**Fig. 3D**].

**Figure 3.**
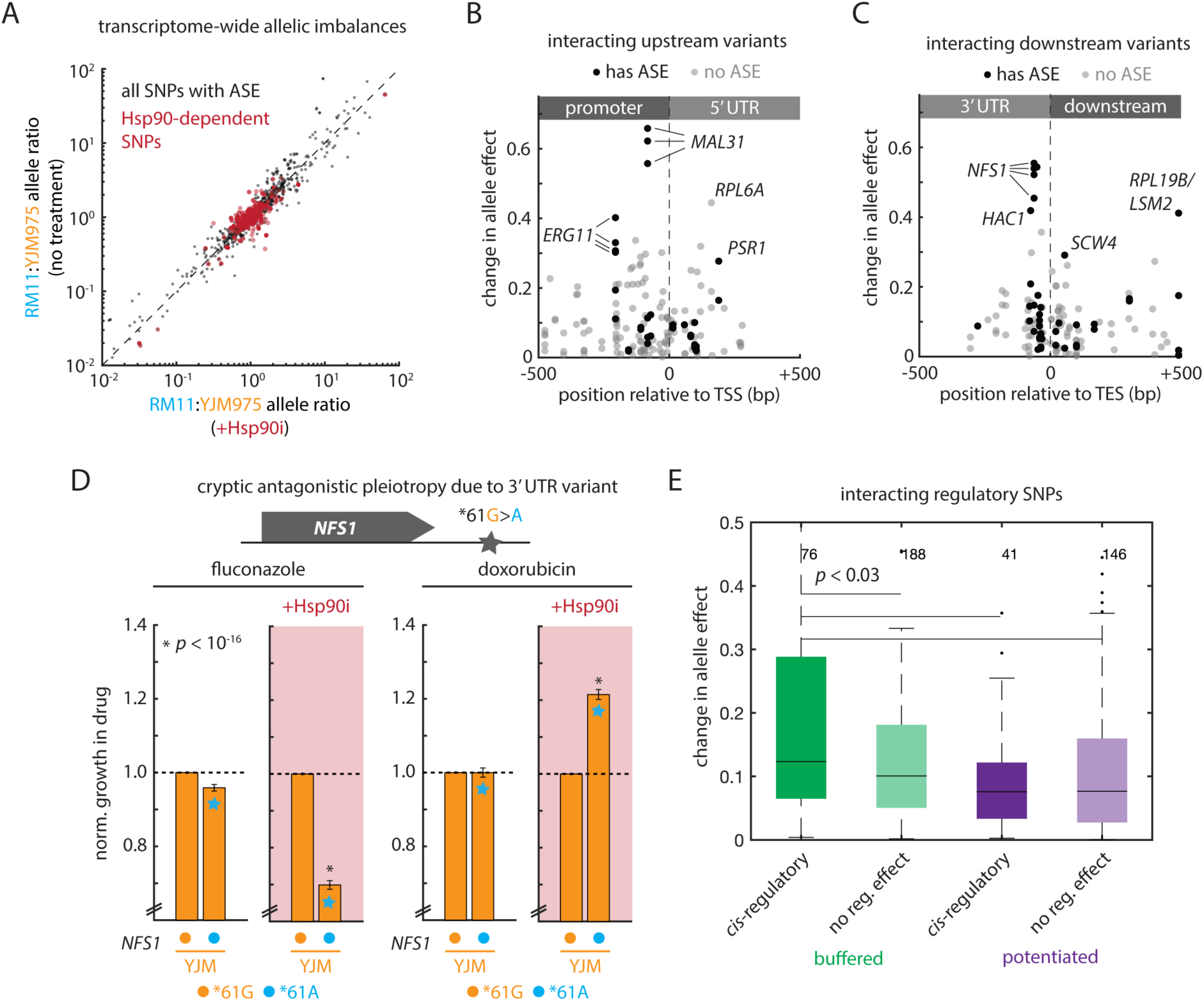
Hsp90 controls the phenotypic outcomes of *cis-*regulatory variation. (A) Allelic ratio (RM counts/YJM975 counts) in the hybrid diploid with Hsp90 inhibited by radicicol (abscissa) or with Hsp90 active (ordinate) for all SNPs with genome-wide significant allele-specific expression (grey) and for SNPs with Hsp90-dependent phenotypic effects (red). Mean allelic ratios are shown across three replicates for each condition. (B) Schematic illustrating the position of upstream Hsp90-modified regulatory variants relative to the TSS of the nearest gene; genome-wide significant allele-specific expression is as indicate (black dots: SNP has ASE [*q* < 0.05]; grey dots: no ASE). Selected large-effect alleles are highlighted. (C) As in (B), but for downstream Hsp90-modified regulatory variants relative to the TES. (D) Growth in fluconazole (left) and doxorubicin (right) of genome-edited strains with Hsp90 active (no shading) and inhibited with radicicol (+Hsp90i; red shading). Each panel shows wild type YJM975 *NFS1*^*61G^, and edited YJM975 *NFS1*^*61A^ (blue stars) strains. Mean and s.e.m. are shown, normalized to the wild type; *p* values by *t* test against the same mutant strain without radicicol treatment; *n* = 384 per genotype. (E) Absolute change in allele effect for Hsp90-modified regulatory variants that were buffered (green) and potentiated (purple), sorted by whether the transcript exhibited allele-specific expression (left) or had a tag SNP but no allele-specific expression (right). Box plots show the median and upper and lower quartiles; whiskers show 1.5 times the interquartile range; *p* values by *t* test; *n* as indicated. See also [**Fig. S3**].

Interestingly, we previously implicated coding variation in Nfs1 in Hsp90-buffered rapamycin resistance arising in a different segregant panel^26^. These observations, in conjunction with the modest effect on ASE, led us to hypothesize that the nature of Hsp90’s impact on genetic variation may be a gene-level property in which the chaperone controls fitness effects by enhancing the amount of folded, active protein that is produced, thereby limiting the phenotypic impact of *cis*-regulatory effects. Indeed, despite having little effect on mRNA ASE itself, Hsp90 more strongly impacted the consequences of potent regulatory variants (polymorphisms associated with significant allelic imbalances; *q* < 0.05) [**Fig. S3D**].

We also noted that more than 200 Hsp90-modified variants were synonymous – that is, they represented a change in codon choice but not in protein sequence. Of these, 38 of 247 (∼ 15%) exhibited mRNA allele-specific expression. As expected, this was a much lower fraction than for regulatory variants, which we expect to act predominantly on mRNA (∼ 29%). Consistent with roles for both mRNA- and protein-level regulation, synonyms with mRNA ASE did not exhibit larger effect sizes than synonymous variants with no ASE [**Fig. S3E**]. Thus, many of the Hsp90-interacting synonymous variants may drive phenotypes via protein levels or co-translational folding. Synonyms without ASE that interacted with Hsp90 indeed exhibited larger changes in normalized translational efficiency (nTE; a statistic that estimates codon optimality based on codon usage and tRNA availability^63^) than Hsp90-dependent synonyms with ASE (which presumably act at the mRNA level) [**Fig. S3E**], further suggesting that protein-level effects may play a role for these variants. On the other hand, synonyms with ASE were closer to protein domain boundaries than those without [**Fig. S3E**], potentially suggesting a role for changes in translation rate and co-translational folding feeding back onto mRNA stability^64^. Thus, synonyms likely constitute an additional form of *cis*-regulatory variant within the open reading frame, analogous to *cis*-regulatory variants outside ORFs that control mRNA expression.

Proteins are typically present at levels that do not limit growth^65^; however, their destabilization upon Hsp90 inhibition might elicit dosage sensitivity, which, in turn, would reveal newfound phenotypes derived from the pre-existing molecular consequences of buffered *cis*-regulatory variants. Consistent with this idea, we found that buffering interactions were stronger than potentiations amongst variants associated with mRNA *cis*-regulation (median 63% larger; *p* < 0.03) [**Fig. 3E**; **Fig. S3F**]. Studies in diverse organisms have illustrated the potential for Hsp90 to have profound and lineage-specific consequences for developmental phenotypes^17^ typically associated with *cis*-regulatory variation, often involving the release of cryptic phenotypic diversity. Our results suggest that these buffering interactions may naturally and frequently arise from the confluence of *cis*-acting variants with the widespread consequences of Hsp90 on the folding and activity of proteins at the core of eukaryotic regulatory circuitry.

### Hsp90 modifies variation in its clients and their targets

Our observation that both coding and non-coding variants drove Hsp90-dependent phenotypes led us to ask whether either were linked directly to Hsp90’s well-characterized chaperone activity [**Fig. 4A**] or instead were connected by complex nonspecific percolation through regulatory circuitry. We initially considered both genetic and physical interactions, as genetic interactions provide information on functional connections that can be challenging to capture physically if the proteins in question are expressed at low levels, or if the relevant interactions are fleeting. Strikingly, modified alleles of all types were much more likely to occur in or regulate genes that interact with Hsp90 (Bonferroni-corrected *q* < 10^-5^ for Hsc82, *q* < 0.001 for Hsp82) [**Fig. 4B**]^66^. We also searched the interactors of Hsp70 chaperones, which collaborate with Hsp90 to catalyze the folding of many client proteins, and those of the Hsp70/Hsp90 Organizing Protein (HOP/Sti1^67–69^). These interactors were enriched for modified alleles (*q* < 10^-5^ for Sti1 and the Hsp70 proteins Ssa1, Ssb1, and Sse1), and the well-characterized Hsp40 adaptor Ydj1 and the kinase co-chaperone Cdc37 exhibited similar enrichments (*q* < 10^-5^) [**Fig. 4B**]. We also noted enrichments amongst the interactors of the Hsp70 homologue Lhs1 (*q* < 0.003) and Hsp40 homologue Xdj1 (*q* < 0.02), suggesting a role for ER and mitochondrial protein quality control in the effects we observed [**Fig. S4AB**]. Although the Hsp90-dependent phenotypes we identified were mechanistically diverse and often driven by *cis*-regulatory variation, most interacting variants (59.8%) occurred in genes that encode direct clients of the Hsp90/Hsp70/Sti1 chaperone system^36^ (*p* < 10^-3^ by Fisher’s exact test). In fact, the genes controlled by interacting *cis*-regulatory variants exhibited particularly strong enrichments for both physical and genetic interactors of Hsp90 and related chaperones [**Fig. 4CD**]. This was consistent with the hypothesis that cells are sensitized to the effects of regulatory mutations on mRNA levels when the client proteins they control are destabilized or otherwise functionally compromised.

**Figure 4.**
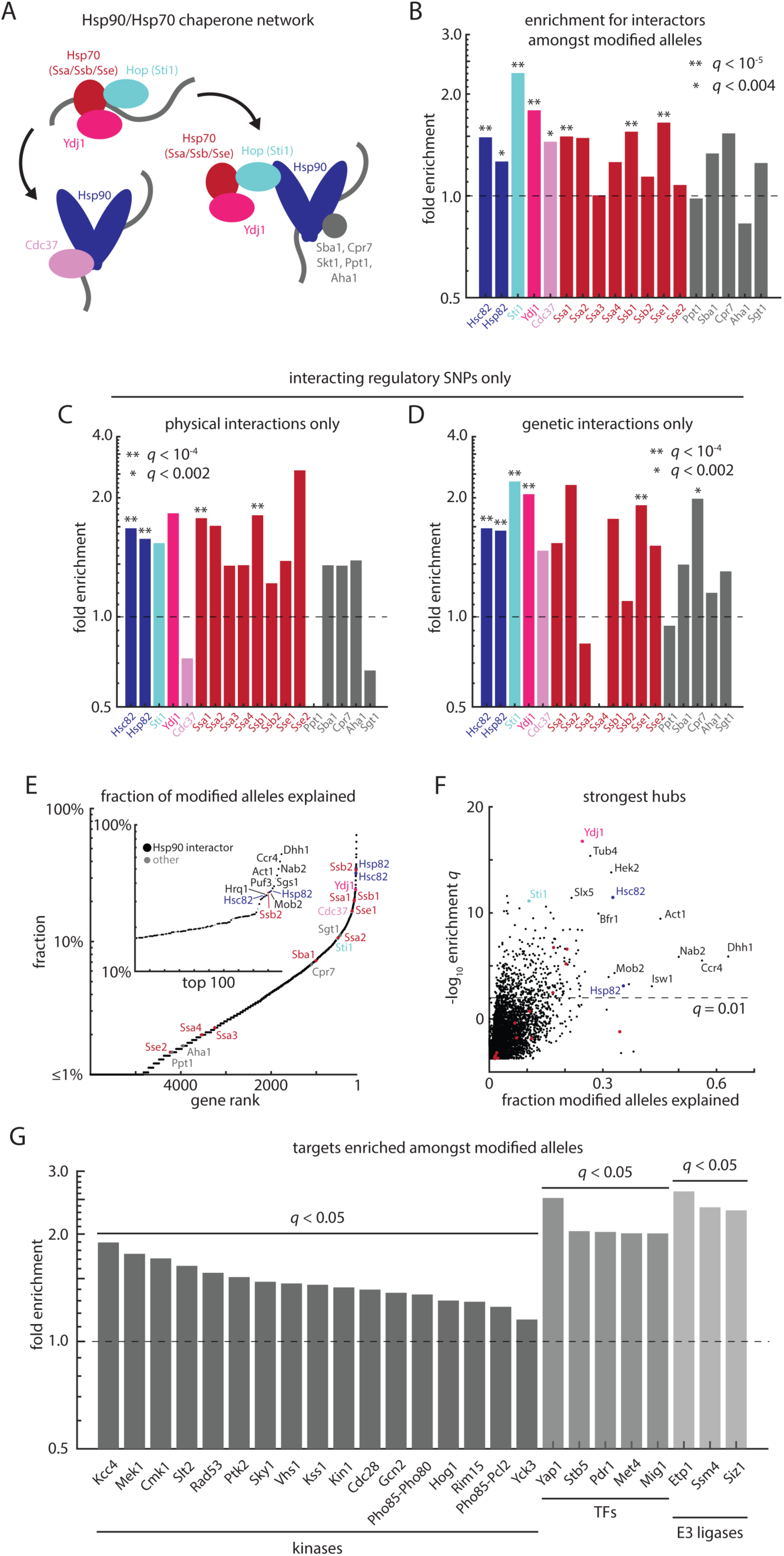
Hsp90-dependent phenotypes arise from the Hsp90/Hsp70 chaperone network and client regulatory proteins. (A) Schematic of the Hsp90/Hsp70 chaperone network and associated co-chaperones and adapter proteins^74^. (B) Relative enrichment of interactors of the indicated proteins amongst Hsp90-modified alleles as compared to all segregating genetic variants. Interactions from the BioGRID database. Bonferroni-corrected *q* values from Fisher’s exact test. (C) As in (B), for only regulatory Hsp90-modified variants and physical interactions from BioGRID. (D) As in (B), for only regulatory Hsp90-modified variants and genetic interactions from BioGRID. (E) Genes ranked by the fraction of their interactors amongst Hsp90-modified alleles. Inset: top 100 hits. Chaperone network components are colored as in (A). (F) Interaction enrichment *q* value (ordinate) as a function of the fraction of interactors amongst Hsp90-modified alleles (abscissa) for all genes in BioGRID. Chaperone network components are colored as in (A). Bonferroni-corrected *q* values from Fisher’s exact test. (G) Relative enrichment of targets of the indicated proteins amongst Hsp90-modified alleles as compared to all segregating genetic variants. Left: kinases; middle: transcription factors; right: E3 ligases. Bonferroni-corrected *q* values from Fisher’s exact test. All proteins shown have *q* < 0.05. See also [**Fig. S4**].

An unbiased analysis to identify the ‘hub’ genes with the most direct biological connections^66^ to the interacting SNPs also highlighted numerous members of the Hsp90/Hsp70 chaperone machinery (*e.g.* Hsc82, Hsp82, Ydj1, Ssa1, Ssb1, Cdc37, Sti1) amongst the genes most connected to the Hsp90-dependent fitness effects we mapped [**Fig. 4EF**]. Moreover, 73 of the top 100 hubs were themselves interactors of Hsc82 or Hsp82 (*p* < 10^-22^, Fisher’s exact test), illustrating the ubiquity of connections from the ‘hub of hubs’ Hsp90 to genes throughout the cellular network. Notably, the hub candidates included not only ubiquitin ligase components such as Slx5 and the kinase activator Mob2, but also numerous RNA-binding proteins including Dhh1, Ccr4, and Nab2 [**Fig. 4F**]. These interactions may point toward multiple mechanisms linking Hsp90 activity to the outcomes of 3’ and 5’ UTR variation, as described above.

To better understand these second-order connections, we examined the targets of three classes of regulatory proteins that are prominent amongst Hsp90’s clientele: kinases, transcription factors (TFs), and E3 ubiquitin ligases^70^. We first assessed which kinases might connect Hsp90 inhibition to changes in the effects of natural variants, identifying seventeen kinases whose targets^38^ were enriched for Hsp90-dependent variation (Bonferroni-corrected *q* value < 0.05) [**Fig. 4G, Fig. S4C**] of which thirteen are known to interact with Hsp90^71^. Two of the remaining four interact with either Hsp70 or the kinase adapter Cdc37. Hsp90 exerted a broad influence via these client kinases: ninety-two predicted targets of the client MAP kinase Slt2, for example, bore interacting alleles^72^. The targets of these enriched kinases accounted for another 32.2% of the interacting alleles in our experiments. Indeed, many of these kinases themselves respond to various stresses (*e.g.*, DNA damage and replication checkpoints [Kcc4, Mek1, Rad53, Cdc28, Pcl2], nutrient response [Gcn2, Pho80/Pho85, Rim15]), heat and cold shock and the UPR [Slt2, Kin1], stress granule regulation [Sky1], and osmotic shock [Hog1]), adding another layer of potential environmental specificity.

Modified variants were also enriched at the targets of five TFs [**Fig. 4G**, **Fig. S4D**; see **Methods**]^39^ but comparatively poor characterization of the substrate specificity of E3 ligases^71^ [see **Methods**] limited our capacity to assess this class of clients, with only three ligases exhibiting significant enrichments [**Fig. 4G**, **Fig. S4E**]^73^. In spite of these limitations, our data establish that > 90% of Hsp90’s effects on the inheritance of biological traits can be explained by genetic variation in direct targets of chaperone function or in the targets of kinases, a deeply conserved class of Hsp90 client proteins. This close coupling enables the simultaneous modification of multiple alleles in complex sub-networks engaged by the integration of diverse stress and signaling inputs that control the function of Hsp90 and its client kinases.

### The influence of Hsp90 on complex heredity

The Hsp90-dependence of phenotype was often highly complex — the median trait comprised 142 modified alleles [**Fig. S5A**] – consistent with a widespread reservoir of co-regulated variants under the influence of Hsp90 activity. Hsp90-dependent growth in maltose, for example, depended on three separate large-effect alleles segregating at the *MAL1*, *MAL3*, and *GAL10/GAL1* loci. The modified loci at *MAL3* and *GAL10* included regulatory variants; the locus at *MAL1* mapped to a deletion encompassing three maltose metabolic genes – *IMA1, MAL11*, and *MAL13*^75^. This segmental deletion was much more deleterious to growth in maltose when Hsp90 was inhibited [**Fig. S4B**], indicating that it was buffered by the chaperone. Gal1 is an Hsp90 client, whereas Mal13, Mal33, and Gal10 are clients of Hsp70^36^ and the kinase Yck3 is thought to act on Ima1^38^, illustrating the far-reaching polygenic impact of the Hsp90 chaperone network.

These joint interactions resulted in a remarkably environment-specific effect of Hsp90 inhibition on fitness [**Fig. 5A**]. Yet despite Hsp90’s pronounced impact in maltose, the three loci had negligible Hsp90 dependence in raffinose [**Fig. 5A**]. This was a general property of Hsp90-dependent heredity in our experiments: the effect of Hsp90 inhibition on the phenotypic consequences of variation was uncorrelated between different stress conditions [**Fig. S5C**]. To better understand how Hsp90 influenced the concerted effects of multiple variants, we examined the eight possible combinations of homozygous genotypes at these three loci and the fitness consequences of evolutionary paths between them. Several of these mutational steps were permitted (did not significantly decrease fitness) exclusively when Hsp90 was either fully active or inhibited [**Fig. S5D**]. This, in turn, altered the mutational paths accessible between genotypes without a stepwise decrease in fitness [**Fig. 5B**]. Simultaneously, Hsp90 inhibition coarsened the landscape: the effect of single-mutation steps^76^ in this mutational neighborhood was increased when Hsp90 was inhibited in maltose, but had little effect on the same transitions in raffinose [**Fig. 5C**]. We observed similar behavior for buffered variants linked to fluconazole resistance at *ERG11*, *PDR1*, and *GRE2* [**Fig. 5DE**], and, as above, these gene products are closely linked to Hsp90 activity: Erg11 is an Hsp90 client, Gre2 a client of Hsp70, and Pdr1 a predicted target of the Rim15, Rad53, Kin1, and Hog1 kinases, among others^36,38^. These results illustrate how Hsp90 can affect polygenic adaptation in ways that are highly selective for a given environment. By altering the effects of mutations from deleterious to permissible (neutral), chaperone activity shapes the tradeoff between metabolic specialization and the accessibility of other phenotypes when the environment changes.

**Figure 5.**
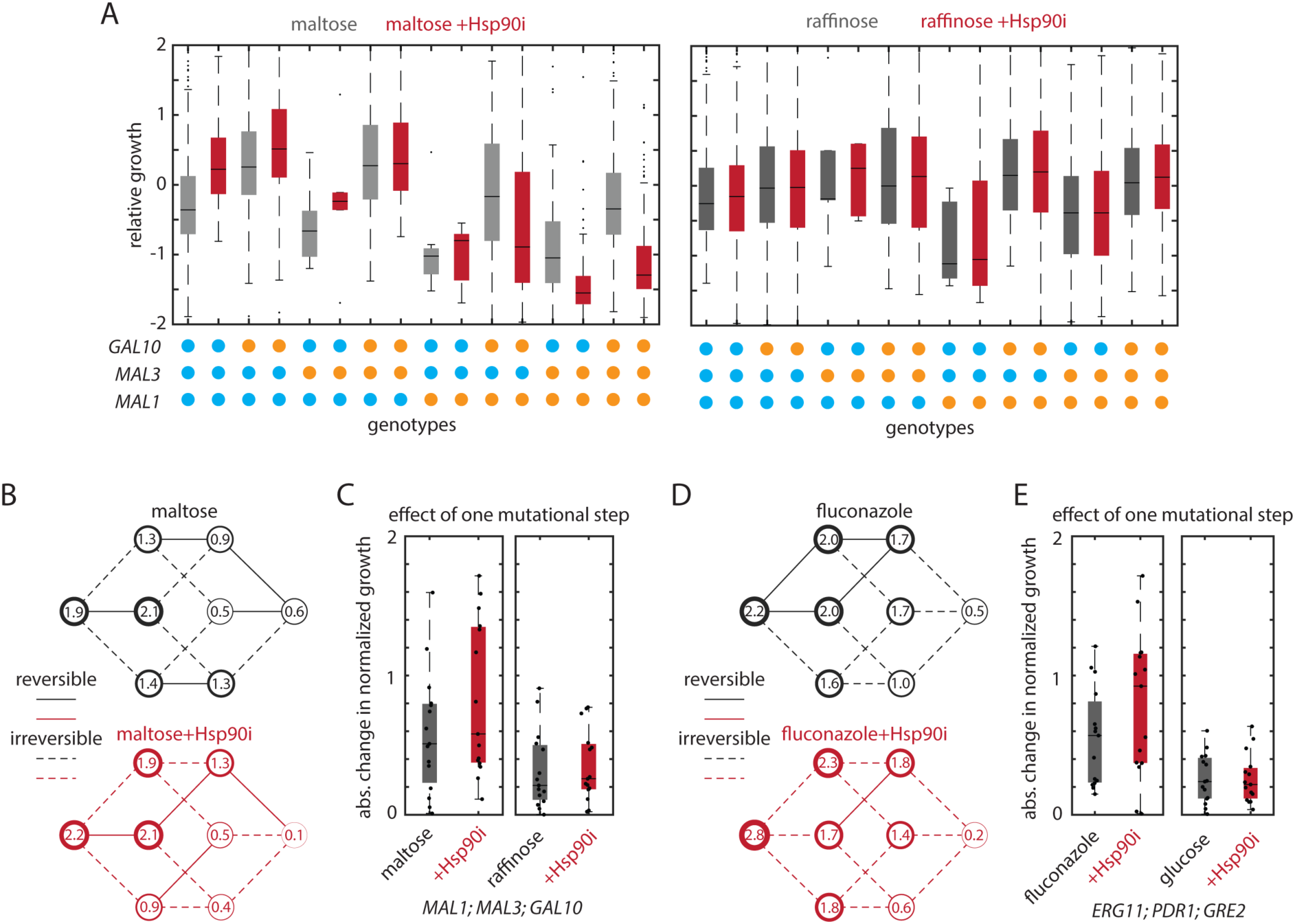
Complex genetic architectures of Hsp90-dependent heredity. (A) Left: Mean normalized growth on maltose of F_6_ segregants binned by their genotypes at *GAL10*, *MAL1*, and *MAL3* and organized by their mutational distance from the fittest genotype (*GAL10^YJM975^*; *MAL1^RM11^*; *MAL3^RM11^*) without (black) and with (red; +Hsp90i) inhibition of Hsp90 by radicicol. Box plots show the median and upper and lower quartiles; whiskers show 1.5 times the interquartile range. Right: As left, but for growth in raffinose. (B) Phenotypic landscape of growth in maltose (top) or maltose with radicicol (bottom). Genotypes at *MAL1, MAL3,* and *GAL10* are as indicated in (A); the fitness of each genotype is encoded in the line width of the enclosing circle. Phenotypic landscapes highlight mutational paths accessible by drift (solid lines; reversible; do not significantly decrease fitness; nominal *p* > 0.05) and paths that alter fitness (dashed lines; irreversible; nominal *p* < 0.05). Indicated within each circle is the fitness of that genotype (in units of standard deviations of F_6_ growth; adjusted such that fitness values are positive). (C) Change in mean normalized growth on maltose for all possible single-mutation steps amongst *GAL10*, *MAL1*, and *MAL3* alleles with (red) without (black) Hsp90 inhibition by radicicol (+Hsp90i). Box plots show the median and upper and lower quartiles; whiskers show 1.5 times the interquartile range. (D) As in (B), but for *ERG11*, *PDR1*, and *GRE2* in fluconazole and glucose. (E) As in (C), but for *ERG11*, *PDR1*, and *GRE2* in fluconazole and glucose. See also [**Fig. S5**].

### Hsp90-dependent variation at young genes

Our identification of interacting variants not only at the *MAL* but also the *FLO* genes, young, rapidly evolving subtelomeric gene families with highly variable gene content across *S. cerevisiae*^77^ [**Supplemental File S2**], prompted us to ask if evolutionarily young genes, as a group, were predisposed to harbor Hsp90-dependent variation. After stratifying all *S. cerevisiae* genes by evolutionary age^78^ [**Fig. 6A**], we found that the youngest genes harbored alleles that exhibited the strongest Hsp90-dependent phenotypic effects [**Fig. 6B**]. Consistent with the prominent role for *cis-*regulation described above, this enrichment was driven predominantly by regulatory variation [**Fig. 6C**; **Fig. S5E**] and strong buffering interactions [**Fig. S5FG**].

**Figure 6.**
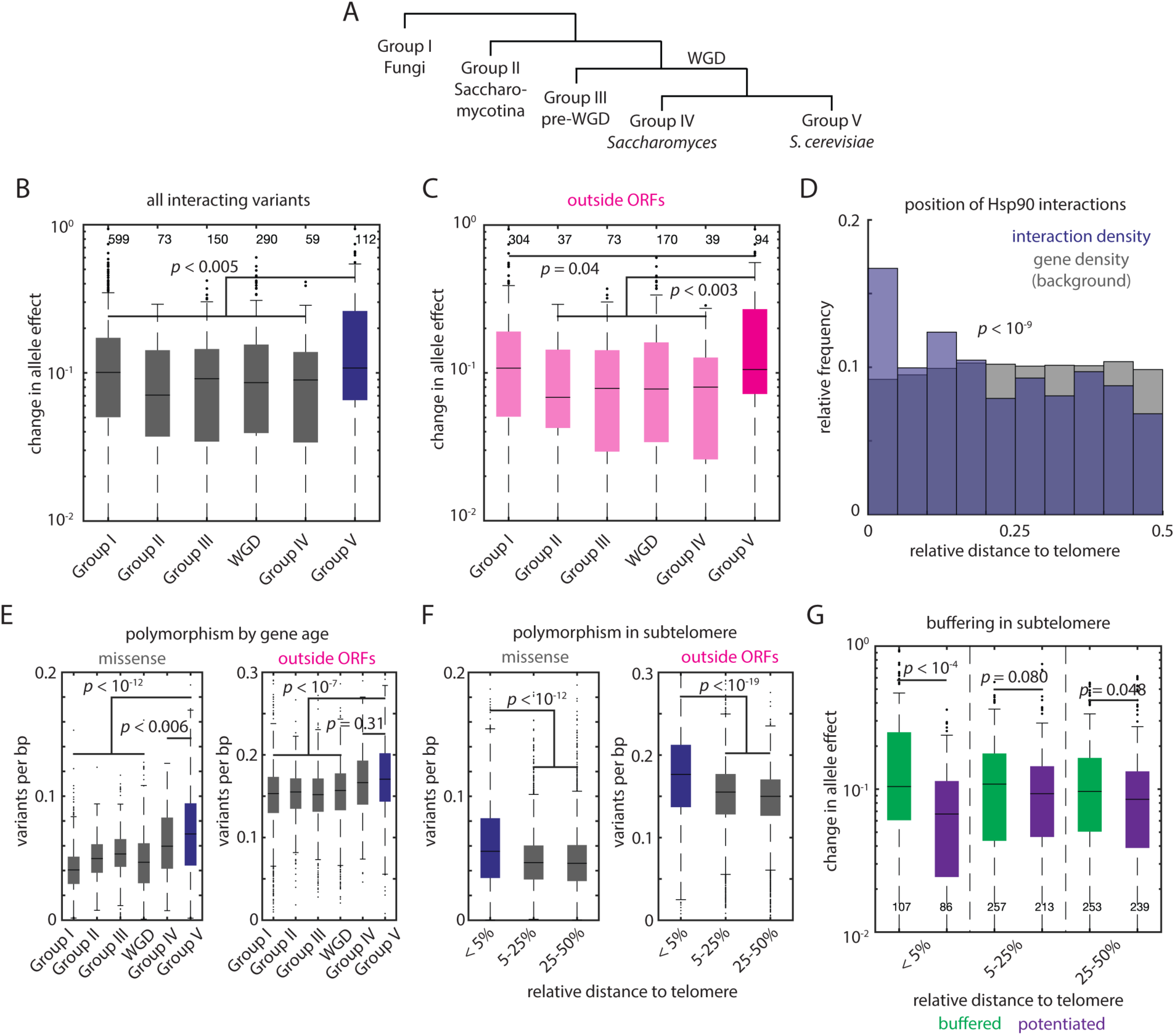
Hsp90 buffers the effects of subtelomeric genetic variation. (A) Schematic of phylogeny used to classify *S. cerevisiae* genes by their evolutionary age based on the presence or absence of homologues in the taxa shown. (B) Absolute change in allele effect for Hsp90-modified variants with ages as indicated. Box plots show the median and upper and lower quartiles; whiskers show 1.5 times the interquartile range; *p* values by Mann-Whitney *U* test. *n* as indicated. (C) As in (B), but only for Hsp90-modified variants outside of open reading frames. (D) Relative frequency of Hsp90-modified variants (blue) and all genes (grey) as a function of distance from the nearest chromosome end (normalized to chromosome length). *p* value by Kolmogorov-Smirnov test. (E) Left: Missense variants per base pair in the 1,002 Yeast Genomes collection for genes with ages as indicated. Right: As left, but for variants outside open reading frames. *p* values by Mann-Whitney *U* test. (F) Left: Missense variants per base pair in the 1,002 Yeast Genomes collection for genes with positions relative to the nearest chromosome end as indicated. Right: As left, but for variants outside open reading frames. *p* values by Mann-Whitney *U* test. (G) Absolute change in allele effect for Hsp90-buffered variants (green) and Hsp90-potentiated variants (purple) with positions relative to the nearest chromosome end as indicated. *p* values by Mann-Whitney *U* test. See also [**Fig. S5**].

In turn, subtelomeric regions across the genome exhibited an unusually high density of Hsp90-dependent mutations relative to their gene content (*p* < 10^-9^) [**Fig. 6D**]. Indeed, even though the set of natural variants we examined harbors an excess of regulatory mutations near chromosome ends, Hsp90-dependent regulatory variants were nonetheless enriched in these hypervariable regions (*p* < 0.03) [**Fig. S5H**]. Across *S.* cerevisiae, both young and subtelomeric genes are enriched for coding and non-coding polymorphisms [**Fig. 6EF**] Moreover, subtelomeric Hsp90-dependent variants exhibited much stronger buffering interactions as compared to potentiations (*p* < 10^-4^) than did variants in the remainder of the chromosome **Fig. 6G**]. Thus, Hsp90-buffered variation provides a potent reservoir of phenotypic novelty^79^, particularly at subtelomeric genes, such as the *MAL* loci, that are known contribute to niche-specific adaptation and harbor more condition-specific alleles of large effect^77^.

### A generalizable model for buffering by Hsp90

How does Hsp90 simultaneously buffer the outcomes of coding and non-coding variation? To investigate this question, we returned to two observations from genetic mapping: first, that Hsp70 and Hsp90’s direct clients were enriched in interacting genes, and second, that regulatory variation, which does not alter protein sequence, was a central driver of Hsp90-dependent heredity. These observations led us to hypothesize that Hsp90-dependence was often an intrinsic property of certain gene products, rather than a property of particular mutations that led one allele to become Hsp90-dependent.

To test this idea, we revisited the buffered *ERG11* regulatory variant [**Fig. 2A**]. Interestingly, genetic mapping also suggested a role for the neighboring Erg11^Asn433Lys^ coding variant in azole resistance when Hsp90 was fully active. We hypothesized that, due to a reduction in Erg11 activity upon Hsp90 inhibition, the effects of both variants would be buffered [**Fig. 7A**]. After reconstructing sensitive strains bearing each variant singly as well as a strain bearing both mutations, we found that, strikingly, the single and double mutants all exhibited buffered resistance to azoles (*p* < 10^-4^; *p* < 0.001) [**Fig. 7B**; **Fig. S6A**].

**Figure 7:**
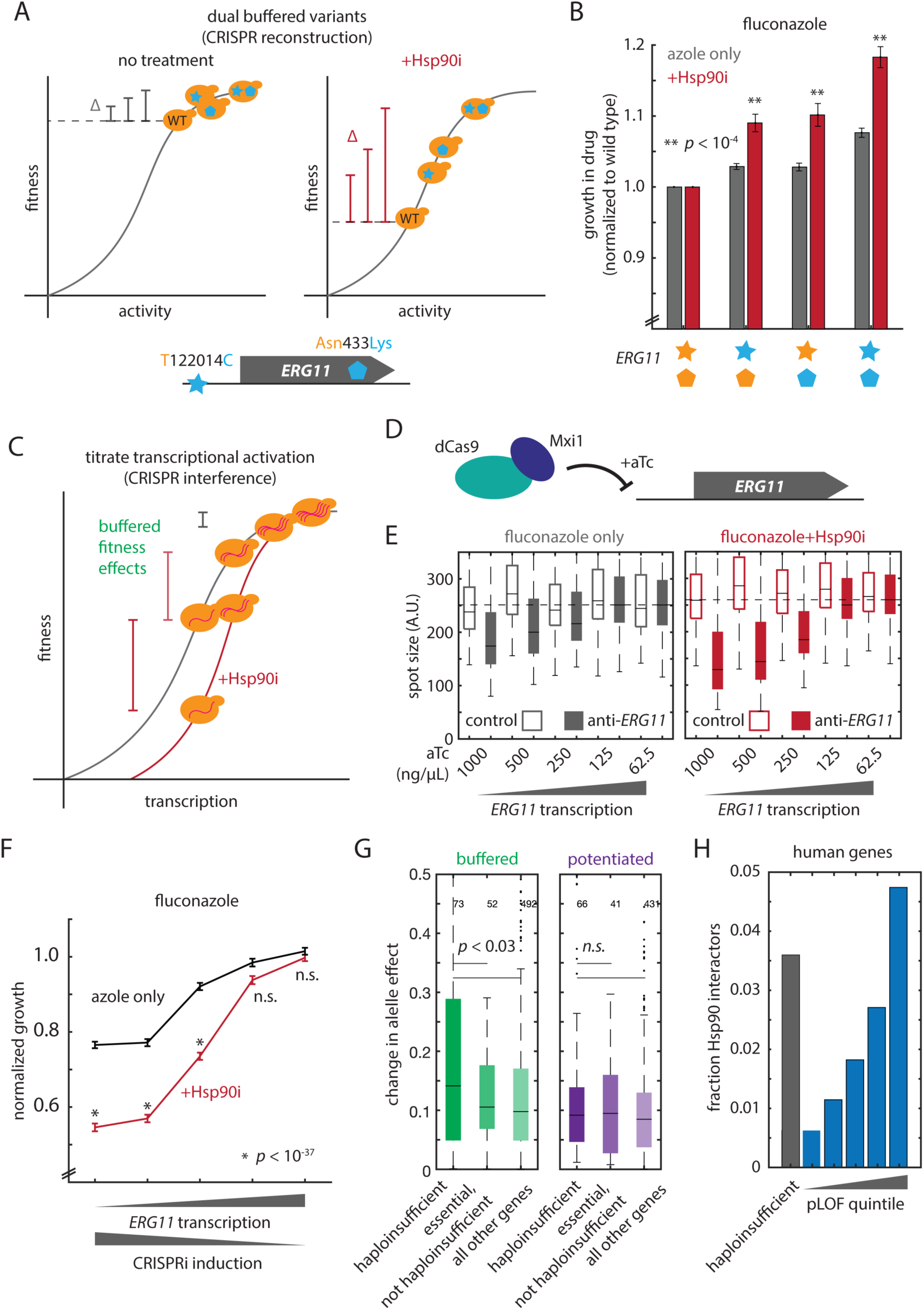
Hsp90 modifies the activity-fitness relationship. (A) Schematic of the effect of Hsp90 inhibition on fitness assuming a non-linear relationship between protein activity and phenotype. Mutations in the hypothetical gene increase activity; the fitness consequences of these changes are amplified when total activity is compromised by Hsp90 inhibition. (B) Growth in fluconazole of genome-edited strains with Hsp90 active (grey) and inhibited with radicicol (+Hsp90i; red). Each panel shows wild type YJM975, edited YJM975 *ERG11*^122014C^, edited YJM975 Erg11^433Lys^, and edited double-mutant YJM975 *ERG11*^122014C^ Erg11^433Lys^ strains. Data shown are mean and s.e.m. normalized to the wild type; *p* values by *t* test against the same mutant strain without radicicol treatment; *n* = 384 per genotype. (C) Schematic of the effect of Hsp90 inhibition on fitness assuming a non-linear relationship between protein activity and phenotype. Increasing CRISPRi expression reduces *ERG11* expression; the fitness consequences of these changes are amplified when total activity is compromised by Hsp90 inhibition. (D) Schematic of CRISPR transcriptional interference mediated by a dCas9:Mxi1 fusion and anhydrotetracycline (+aTc) inducible gRNA. (E) Growth of YJM975 haploid strains expressing either a non-targeting control gRNA (open boxes) or an anti-*ERG11* gRNA (closed boxes) in fluconazole without the addition of radicicol (grey boxes) or with radicicol (red boxes). Shown is the concentration of inducing molecule (aTc) added to each culture. Box plots show the median and upper and lower quartiles; whiskers show 1.5 times the interquartile range; *n* = 384 per genotype. (F) Data in (E) normalized to the growth of the YJM975 strain expressing the non-targeting control gRNA. Growth is in fluconazole without the addition of radicicol (black) or with radicicol (+Hsp90i; red). The concentrations of inducing molecule (aTc) added to each culture are (left to right) 1000 ng/µL, 500 ng/µL, 250 ng/µL, 125 ng/µL, and 62.5 ng/µL. Data shown are mean and s.e.m. (G) Absolute change in allele effect for Hsp90-modified variants in haploinsufficient genes, essential but not haploinsufficient genes, or all other genes, as indicated. Buffered interactions are shown at left (green) and potentiated interactions at right (purple). Box plots show the median and upper and lower quartiles; whiskers show 1.5 times the interquartile range; *p* values by *t* test; *n* as indicated. (H) Fraction of Hsp90-interacting gene products amongst all human genes, divided in quintiles by increasing gnomAD pLOF score (blue) and amongst genes haploinsufficient in embryonic stem cells (grey). See also [**Fig. S6**].

We next sought to explain why these distinct mutations were affected similarly by Hsp90. Building on our observation that *cis*-regulatory variation is frequently buffered, we hypothesized that this behavior could arise if activity-to-fitness relationships for a given gene modified by the chaperone were sigmoidal rather than linear [**Fig. 7C**]. Sigmoidal relationships of this kind have been observed empirically for combinations of mutations in proteins^80,81^ and for combinations of *cis*-regulatory variants at different genes^82^. Under normal physiological conditions, most such relationships are relatively flat, permitting modestly perturbative variants to accumulate. However, when environmental stress depletes the reservoir of Hsp90 function, sigmoidal activity-to-fitness relationships would be transformed for genes that interact with the chaperone or its client kinases, shifting toward the precipice of a high slope regime. The previously tolerated molecular consequences of variants that cause the protein to be improperly folded, aberrantly post-translationally modified, or decrease its expression would have a very different fitness impact due to the increased local slope of the activity-fitness sigmoid, amplifying the phenotypic impact of buffered variants. Such a model has a striking prediction: for a given locus such as *ERG11,* the fitness effects of arbitrary mutations should depend on Hsp90 activity, and the phenotypic consequences of reducing Erg11 activity should be buffered.

To test these predictions quantitatively, we used synthetic CRISPR interference^83^ to reduce *ERG11* transcription in the YJM975 parent [**Fig. 7D**]. As compared to a non-targeting guide RNA, reducing *ERG11* levels [**Fig. S6B**] caused a dose-dependent sensitivity to fluconazole [**Fig. 7E**]. In turn, Hsp90 inhibition caused a progressively greater sensitivity concomitant with decreasing *ERG11* transcription (*p* < 10^-8^) [**Fig. 7EF**]. That is, the phenotypic consequences of reduced Erg11 activity were systematically buffered by Hsp90, just as we had observed for natural variation. Reducing *ERG11* levels had no effect in the absence of azole, regardless of Hsp90 activity [**Fig. S6C**], and comparable results were obtained in the RM11 parent, indicating that the phenomenon was a general property of reduced Erg11 levels rather than a property of a specific *ERG11* genotype [**Fig. S6D**].

Finally, we asked whether we could identify genes with a steep activity-fitness relationship that might be more prone to Hsp90-dependent buffering. We examined haploinsufficient genes—those whose heterozygous mutants affect the diploid—because reducing their expression by approximately one-half leads to an appreciable phenotype [**Fig. S6E**]. Indeed, mutations in these genes^84^ had stronger Hsp90-dependent effects [**Fig. S6F**]. This did not arise simply because many haploinsufficient genes are essential: essential genes that are not haploinsufficient did not exhibit this trend. Moreover, in concordance with our model, these enrichments arose exclusively amongst buffering interactions [**Fig. 7G**]. Intriguingly, human genes scored as more essential based on the frequency of predicted loss-of-function mutants in gnomAD^85^ or identified as haploinsufficient in experiments in embryonic stem cells^86^ are also enriched for known Hsp90 interactors^87^ [**Fig. 7H**]. These genes may likewise be candidates to harbor Hsp90-buffered genetic variation arising from sigmoidal activity-fitness relationships.

## DISCUSSION

The influence of Hsp90 on the developmental and adaptive consequences of genetic variation has long been appreciated. However, aside from a handful of examples of mutations in client proteins, the molecular underpinnings of these effects have remained enigmatic. Here, the unprecedented scale of our nucleotide-resolution approach revealed a surprise: *cis-*regulatory variants, which by definition do not alter protein stability, emerged as a driver of nearly half of Hsp90-dependent phenotypes. This behavior arises naturally from the chaperone’s pervasive influence on the activity-fitness relationships of core regulatory proteins. Buffering interactions, which are enriched amongst these relationships, position Hsp90 to govern the release of cryptic, potentially adaptive variation in this regulatory circuitry. This mechanism combines the privileged evolutionary role of *cis*-regulatory variation^88^, which enables sophisticated spatiotemporal programming, with the exquisite stress-dependent regulation of the Hsp90 chaperone machinery to release adaptive phenotypes under stress.

The preponderance of this behavior in the concise budding yeast genome suggests that interactions between Hsp90 and regulatory variation may be even more prominent in organisms with more elaborate regulatory architectures. Considering the modest number of conditions we surveyed in our experiments, the involvement of more than 10% of genes in Hsp90-dependent phenotypes emphasizes the widespread influence of Hsp90 on heredity. Indeed, the pragmatic need for a low density of polymorphisms in our panel means that our results likely represent a lower bound on the prevalence of this phenomenon, even in budding yeast. Significant technical advances are still needed to understand Hsp90-dependent alleles of small effect and genetic epistasis that is itself controlled by chaperone activity, both of which may play a role in Hsp90-dependent heredity. Nonetheless, even generous estimates of the contribution of epistasis suggest that, for some phenotypic transformations driven by the chaperone, other heritable variation that depends on Hsp90 activity (*e.g.*, prions^89^ or long-lived chromatin modifications^6^) may also play an important role.

Our experiments identified numerous instances in which the effect of Hsp90 activity on fixed natural polymorphisms was as large as the effect of the variant alone. These were widespread throughout the genome, and although it is not straightforward to infer selection coefficients directly from genetic mapping experiments, the effects we observed likely exceed the threshold for effective selection by multiple orders of magnitude. That many Hsp90-contingent effects were buffered and highly condition-specific further suggests a role in adaptation. Connecting a wide array of mutations to chaperone activity both directly and *via* client regulatory proteins may confer a particular advantage in rapidly changing environments^90^, when modular regulons can be engaged in a stress-dependent fashion (*e.g.* by kinases).

Indeed, multiple mechanisms may converge on a single protein: many interacting alleles at direct clients were also *trans* targets of client regulators. The environmental specificity of these interactions means that buffered variation can accumulate under normal conditions without compromising fitness. At the same time, this variation is likely shaped by repeated selection during periods when Hsp90 activity is compromised. This allows a greater diversity of genetic variation to persist in the population^91^, broadening the accessible adaptive trajectories under stress. Moreover, by virtue of repeated cycles of selection, this reservoir of genetic potential may be predisposed to drive adaptive phenotypes^92^. Our results also explain why Hsp90 has such a profound impact on development, where the revelation of phenotypes due to *cis*-acting regulation would have especially potent consequences.

Might there be an advantage to maintaining such a dependence on Hsp90? Some connections may be an inevitable consequence of kinase structure and dynamics, although the diversity of client status within kinases families suggests that these proteins can readily evolve away from chaperone dependence^8^. Our results illustrate a potential evolutionary advantage for the retention of this dependence. Sigmoidal activity-to-fitness relationships, which are common among Hsp90 clients, limit the fitness effects of modest perturbations to a protein’s activity. Yet they also amplify the phenotypic consequences of variants that spark larger excursions in activity. Coupling Hsp90 function to the shape of such activity-to-fitness relationships thus provides a route for organisms to reconcile the competing evolutionary demands of robustness and evolvability by modulating the fitness consequences of changes in activity in response to environmental conditions.

The mechanisms we have discovered allow the evolutionary potential of variants of large effect to be harnessed precisely when organisms are most ill-suited to their environments. The involvement of highly polymorphic subtelomeric genes may enhance this phenomenon, as may other environmentally responsive mechanisms, such as the Hsp90-dependent [*ESI^+^*] prion. This epigenetic element regulates the expression of previously silent genetic information in precisely these regions^93^. In some instances, Hsp90 contingency may even be co-opted as a regulatory strategy. Haploinsufficient genes, which are enriched in Hsp90 clients, may preferentially harbor such buffered *cis*-regulatory effects, and our data suggest *cis*-regulatory variants controlling proteins known to interact with Hsp90 or its client kinases may be especially profitable to explore. Understanding how the integration of Hsp90 and kinase signaling modules enable re-interpretation of the genotype-to-phenotype relationship in different environmental contexts is also an exciting avenue for future discovery.

Prior studies have indicated that Hsp90 facilitates the accumulation of coding variation in clients^94^, accelerating their evolution^95^. Our findings suggest that the chaperone enables the accumulation of cryptic regulatory variation at these genes at least as strongly, which provides an unappreciated and appealing potential mechanism for the influence of Hsp90 on developmental evolution in diverse systems from fruit flies to Mexican cavefish. Likewise, in cancer and in pathogens, the buffering of drug-resistant regulatory mutations by Hsp90 may allow their accumulation, subsequently being enriched amongst the surviving cells following treatment and promoting future drug-resistance. This, in turn, suggests that a paradigm favoring therapeutic Hsp90 inhibition, predicated on the Hsp90-addicted v-Src kinase, may be counterproductive if cryptic, buffered resistance is revealed and enriched. Distinguishing cases in which Hsp90 inhibition is helpful or deleterious to treatment is an important question for future investigation, as is the relationship between an individual’s reservoir of Hsp90-dependent buffering capacity and their polygenic risk for genetic disease. The decline of this buffering reservoir throughout the lifespan may also contribute to the emergence of pathologies with age.

Our findings address long-standing mechanistic questions surrounding the role of Hsp90 in transmuting genetic variation into selectable phenotypes. Hsp90’s activity and connectivity with an expansive regulon of client proteins position it to control activity-fitness relationships, and their evolution, throughout the core eukaryotic regulatory circuitry. In turn, the effects of numerous coding and regulatory variants simultaneously depend strongly on its stress-regulated activity. Finally, the acute environmental specificity of these buffered fitness effects facilitates their accumulation in advance of stress. Together, these features render Hsp90 a central governor of the critical evolutionary tradeoff between robustness and adaptability in the face of a changing environment, when diversification is most crucial.

## Acknowledgments

We thank Joseph Schacherer for the 1,002 Yeast Genomes Project strain collection, Justin Smith for the CRISPR interference plasmid, and the Jarosz Lab for critical feedback and advice. This work was supported by the NIH (DP2-GM119140, RF1-AG057334, R01-AG06341801, and R01-HG012366 to D.F.J.; F32-GM125162 to C.M.J.), the National Science Foundation (NSF-MCB116762 to D.F.J.), the Swiss National Science Foundation (P2ZHP3_174735 to J.A.-R.), the European Molecular Biology Organization (ALTF 724-2018 to J.A.-R.), a Searle Scholar Award (14-SSP-210 to D.F.J.), a Kimmel Scholar Award (SKF-15-154 to D.F.J.), a Vallee Scholar Award (to D.F.J.) and a Discovery Innovation Award from Stanford University (to D.F.J.). D.F.J. is also a Science and Engineering Fellow of the David and Lucile Packard Foundation. Some of the computing for this project was performed on the Sherlock cluster. We would like to thank Stanford University and the Stanford Research Computing Center for providing computational resources and support that contributed to these research results.

## Author contributions

C.M.J., J.A.-R., and D.F.J. conceived the project. C.M.J. performed pilot, mapping, genome editing, phenotyping, and transcriptomic experiments, designed and implemented the genetic mapping pipeline, performed transcriptomic and bioinformatic analyses, and prepared the figures. J.A.-R. performed pilot experiments, designed genome editing reagents, and performed genome editing experiments. C.M.J. wrote the original draft of the manuscript. C.M.J. and D.F.J. reviewed and edited the manuscript with input from J.A.-R.

## List of Supplemental Figures and Tables

Supplemental Figure S1: Related to Figure 1.

Supplemental Figure S2: Related to Figure 2.

Supplemental Figure S3: Related to Figure 3.

Supplemental Figure S4: Related to Figure 4.

Supplemental Figure S5: Related to Figure 5 and Figure 6.

Supplemental Figure S6: Related to Figure 7.

Supplemental Figure S7: Related to Method Details.

Supplemental Table S1: Stress conditions used in this study.

## STAR METHODS

### Resource availability

#### Lead contact

Further information and requests for resources and reagents should be directed to and will be fulfilled by Prof. Daniel F. Jarosz.

#### Materials availability

The original F_6_ haploid genetic mapping panel is available from the National Collection of Yeast Cultures and the F_6_ diploid mapping panel and all other yeast strains described here are available upon request to Prof. Daniel F. Jarosz.Bacterial strains bearing plasmids used here are likewise available upon request. Various CRISPEY gene editing plasmids are available from AddGene.

#### Data and code availability

All data supporting the findings of this study are available within the paper and its Supplemental Materials. All data required to recapitulate the mapping results and analyses are available at https://www.dropbox.com/scl/fo/0bx04fxbmzdm2tx63o2im/ACqBYF9kjZCe0DyLybhnJzU?rlke y=49cj84huufa5b2j7ikvamgigf&dl=0. All code required to generate the figures is available at https://github.com/cjakobson/hsp90-mapping. Upon publication a snapshot of the code and dependencies will be deposited as a Zenodo repository. RNA sequencing data are available at the Gene Expression Omnibus (GSE242925). Raw images of phenotyping experiments and custom code for cropping images and estimating spot sizes are available upon reasonable request to Prof. Daniel F. Jarosz.

### Experimental model details

#### Yeast strains

The construction and genotypes of the parents of the F_6_ haploid collection, RM11 and YJM975, are described in She and Jarosz^31^. Briefly, the F_6_ diploids used here were constructed by mating *leu*^-^*HYG*^R^ and *LEU*^+^*hyg*^S^ F_6_ haploid isolates genotyped previously^31^ on YPD agar [RPI] followed by selection for diploidy on synthetic defined medium lacking leucine [Sunrise] and supplemented with 200 mg/L hygromycin. These F_6_ diploids were arrayed into 384-well microtiter plates for cryostorage in SD-Leu+Hyg with 15% glycerol. All robotic manipulations were carried out using a Singer ROTOR pinning robot. The RM11 and YJM975 haploid parents of the original genetic cross bear auxotrophies used to facilitate crossing that are not present in the final F_6_ diploid mapping panel. To avoid undesired interactions with these auxotrophies during validation experiments, reconstructions of Hsp90-interacting alleles were carried out in strains YDJ6635 (YJM975a from the SGRP haploid collection^32^) and YDJ6649 (a spore derived from diploid RM11a/α), both of which are auxotrophic only for uracil. Key yeast strains and plasmids are listed in the **Key Resources Table**.

#### Yeast culture conditions

Unless otherwise noted, yeast were propagated and phenotyped in minimal glucose medium consisting of 6.7 g/L yeast nitrogen base [RPI] and 20 g/L glucose [Sigma] supplemented with 20 mg/L uracil [Sigma] to satisfy the lone auxotrophy of the F_6_ diploid panel. Prior to phenotyping experiments, diploid strains were revived from frozen stocks in liquid SD-Leu+Hyg medium in 384-well microtiter plates, propagated 24 hr at 30°C, then pinned to 1536-spot arrays on minimal glucose agar PlusPlates [Singer] and grown for 24 hr – 48 hr at 30°C. 1536-spot source plates prepared as described above were replicated to conditions as indicated using a Singer ROTOR pinning robot. Plates were incubated at 30°C and imaged at 24 hr intervals using an Epson V800 scanner. Images were cropped using custom R scripts and spot sizes were quantified using the *gitter* R package. Colony sizes were extracted for genetic mapping using custom MATLAB scripts. Working concentrations for the various environmental perturbations are described in **Supplemental Table S1**.

### Method details

#### *v-Src* assay for Hsp90 activity

*v-Src* was expressed under the control of a galactose-inducible promoter from PDJ308 in the F_0_ heterozygous diploid YDJ5924. Cells were grown on solid medium containing either 2% glucose (repressing) or 2% galactose (inducing) as the sole carbon source and with or without 10 µM radicicol; Images were cropped using custom R scripts and spot sizes were quantified using the *gitter* R package^96^.

#### Western blotting

Cell lysis was conducted by bead beating in 20% trichloroacetic acid for 10 min at 50 Hz. Cell mass was normalized by optical density prior to lysis; approximately 1 OD-mL of cell mass was used per sample. Protein was precipitated for 30 min at ∼23,000 x g at 4°C and the precipitate was resuspended in Tris pH 7.5 with 2% SDS before denaturation at 95°C for 5 min. Polyacrylamide gel electrophoresis was performed using 12-20% gradient gels at 120 V for 75 min. Proteins were transferred to PVDF membranes in Towbin transfer buffer for 2 hr at 100 V. Membranes were blocked with rehydrated milk, probed overnight at 4°C, and developed using WestFemto chemiluminescent substrate. Matched gels loaded with equal sample volumes were stained with Coomassie brilliant blue in parallel to western blotting. When required for reprobing, membranes were stripped of primary and secondary antibodies by boiling in Tris-buffered saline for 5 min. Antibodies are listed below.

#### Genome editing using CRISPEY

Oligonucleotides containing appropriate guide and donor sequences for CRISPEY editing were designed as described by Sharon and colleagues^35^ using code provided in that study. Oligos were designed separately for RM11 to YJM975 and YJM975 to RM11 mutations to account for differences in guide and template sequences due to neighboring polymorphisms (see **Key Resources Table**). SRR800790 and SRR1569760 were used as reference genomes for YJM975 and RM11, respectively. Gene editing was conducted largely as described in Sharon and colleagues^35^. Briefly, dsDNA was synthesized from oligos by PCR, dsDNA fragments were inserted into the NotI-digested PDJ2318 editing plasmid by isothermal assembly using a 2xHiFi Assembly kit, and the ligated DNA was transformed into chemically competent *E. coli*. Gene editing of yeast was performed by transformation of the editing plasmid and selection on SD-Ura followed by 24h of growth in synthetic galactose medium lacking uracil. From these editing reactions, yeast cells were streaked to single colonies on YPD plates followed by restreaking to 5-FOA plates to cure the editing plasmid. Single colonies were genotyped by PCR and Sanger sequencing of the desired edited locus; screening two colonies per edit was typically sufficient to recover an edited clone.

#### Homozygous diploid mRNA sequencing

Cell were grown in triplicate in minimal media as described above with and without 10 µM radicicol and harvested at OD ∼ 0.8-1.0. Cell pellets were snap-frozen and stored at −80°C before submission to BGI for RNA extraction, polyA enrichment, reverse transcription, and sequencing, as described above for allele-specific expression analysis. Transcript abundance was again estimated using *kallisto*. Transcripts per million (TPM) estimates agreed well between replicates [**Fig. S7**]. Results are summarized in **Supplemental Table S4**.

#### CRISPR interference

CRISPRi plasmids were cloned as described in Smith *et al.* Briefly, the anti-*ERG11* guide RNA sequence was ordered as an ssDNA oligonucleotide and amplified using extension primers as described in^83^. The CRISPRi plasmid (p416gT-Gal4-dCas9-Mxi1) was opened with NotI and the anti-*ERG11* guide inserted by isothermal assembly using a 2xHiFi Assembly kit, and the ligated DNA was transformed into chemically competent *E. coli* to create PDJ2494. After transformation into *S. cerevisiae* alongside a matched control plasmid with a non-targeting gRNA (PDJ397), strains were pre-grown for 24 hrs in the appropriate concentration of anhydrotetracycline (aTc) to establish steady-state knock-down then subjected to phenotyping under the appropriate aTc concentrations as described. Knock-down of Erg11 was quantified using a C-terminal fusion of mNeonGreen to the *ERG11* ORF^97^.

### Quantification and statistical analysis

#### Genetic mapping

Prior to analysis, colony sizes were normalized to mean 0 and variance 1 (*i.e.* Z-scored) and filtered for outliers attributable to missing spots or debris (due to occasional pinning errors or image artifacts). Trait vectors were filtered for strains with an excess of missing genotypes and markers with insufficient genotype coverage were excluded. Statistical genetic mapping was performed essentially as described previously^30^, with the difference that in the present work we mapped the change in the Z-scored trait for each segregant in each condition with and without 10 µM radicicol treatment. Briefly, coarse mapping was conducted by sequential stepwise selection on the homozygous followed by full (homozygous and heterozygous) genotype matrices. Fine mapping was conducted by ANOVA on *in silico* allele swaps as described in She and Jarosz 2018^31^. The genotype matrix included geometric pseudogenotypes corresponding to edge wells and to capture systematic plate-to-plate variability. Loci that were fine-mapped for one trait were inferred to be the underlying causal variant for other traits in which they were the lead candidate.

#### 1002 Genomes yeast strains and phylogenetic analysis

Genotype information for the 1,002 Genomes collection and the corresponding yeast strains^33^ were a generous gift of Joseph Schacherer (Univ. of Strasbourg). Frozen stocks were revived and phenotyping experiments performed as described above for the F_6_ diploid mapping panel. Growth measurements and quantification were conducted by optical scanning of Singer PlusPlates using *gitter* and custom code as described above.

#### Allele-specific expression analysis

Cells were grown in triplicate in minimal media as described above with and without 10 µM radicicol and harvested at OD ∼ 0.8-1.0. Cell pellets were snap-frozen and stored at −80°C before submission to BGI (www.bgi.com/us/home) for RNA extraction, polyA enrichment, reverse transcription, and sequencing. Transcript abundance was estimated using *kallisto*^98^ and read counts per-site were estimated with *bcftools*^99^ from *kallisto* pseudo-alignments. Allele-specific expression was called per-site based on a binomial test against the prior of major allele frequency = 0.5, with genome-wide Bonferroni correction based on the number of transcribed biallelic sites. Indels were excluded. After per-site calling, SNPs were assigned to genes and genes were assigned allelic imbalances if any SNP associated with their transcript exhibited significant allele-specific expression in any of the three replicate sequencing samples. Allelic imbalances agreed well between replicates [**Fig. S7**]. Results are summarized in **Supplemental Table S3**. Sensitivity estimates were conducted based on a sliding simulated read depth followed by binomial tests, with mock Bonferroni correction performed assuming the true number of transcribed biallelic sites in the hybrid diploid.

#### Bioinformatic resources and enrichment analyses

Genomic analyses used the *S. cerevisiae* R64 reference genome unless otherwise noted. Transcription start and end sites were retrieved from^100^. Targets of sequence-specific transcription factors were retrieved from^39^. Predicted kinase targets were retrieved from^38^. Enriched TF hits were logically connected to Hsp90 function: Mig1^101^ and Pdr1^25^ interact with Hsp90; Hsp90 inhibition downregulates Met4 targets^7^; and Yap1^102^ and Stb5^103^ homologues interact with Hsp90 in other species. Modified alleles at targets of Pdr1 and Stb5 were enriched for fitness effects in fluconazole, consistent with the role of these TFs in multidrug resistance. Although several E3 ligases exhibited modest enrichments for BioGRID physical interactors, only three (Etp1, Siz1, and Ssm4) were significant (*q* < 0.05) after multiple-testing correction. We also examined the targets of RNA-binding proteins based on PAR-CLIP and other interaction data (from the POSTAR3 database), but did not observe any significant enrichments (*q* < 0.05) after multiple-testing correction^104^. Gene and protein descriptions and GO terms were retrieved from SGD^71^ and UniProt^105^. Predicted protein structures were retrieved from the AlphaFold Protein Structure Database^51^. Gene age estimates were retrieved from^78^. Haploinsufficient genes in *S. cerevisiae* were retrieved from^84^. pLoF data was retrieved from^85^. Haploinsufficient genes in human embryonic stem cells were retrieved from^86^.

### Key Resources Table

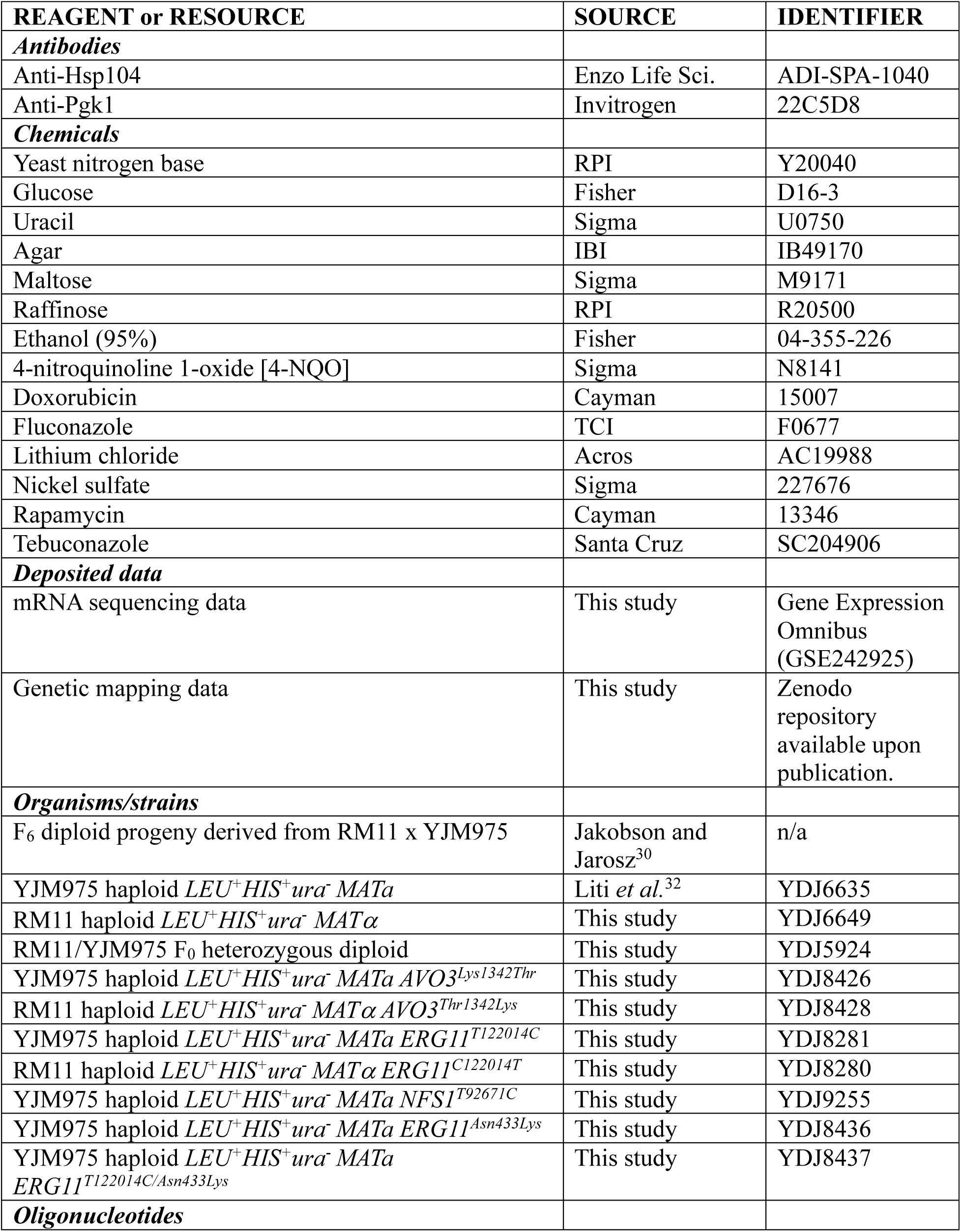
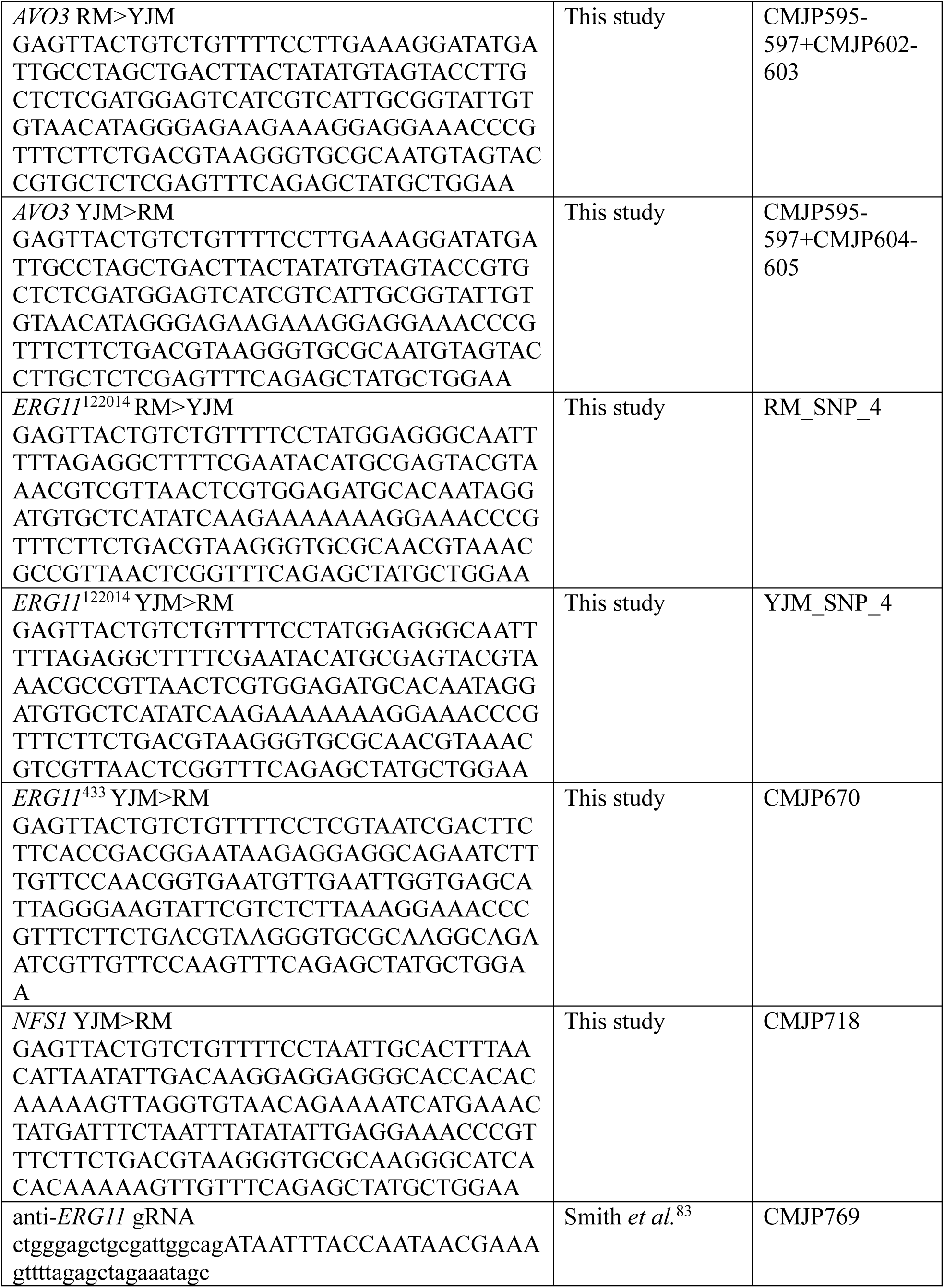
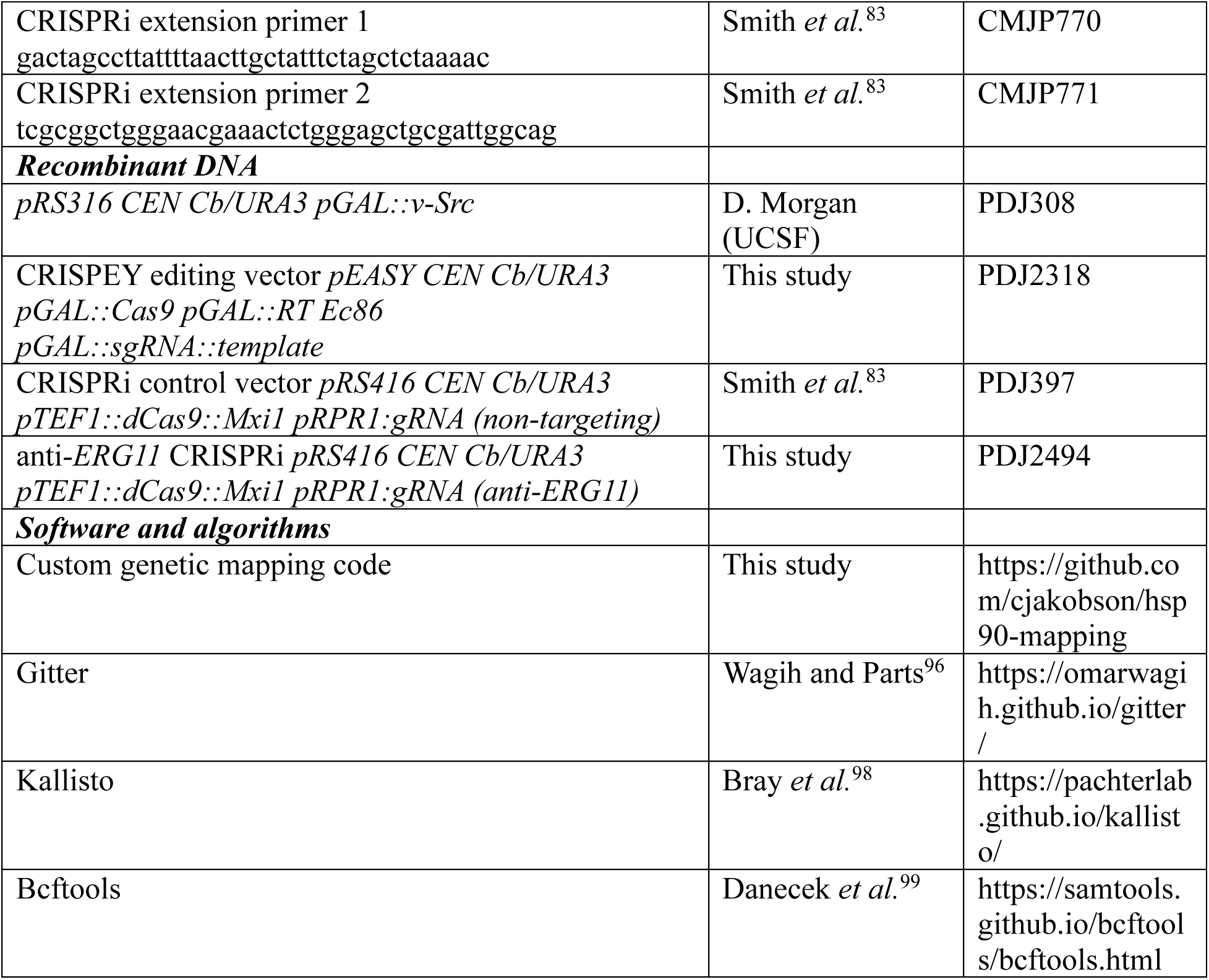

## List of Other Supplemental Items

Supplemental Table S2: Summary of genetic mapping results.

Supplemental Table S3: Allele-specific expression data.

Supplemental Table S4: Homozygous diploid mRNA expression data.

