## Supplemental Information for "Regulatory variants drive Hsp90’s effect on adaptation"

**Supplementary Information**

To accompany Jakobson *et al.*

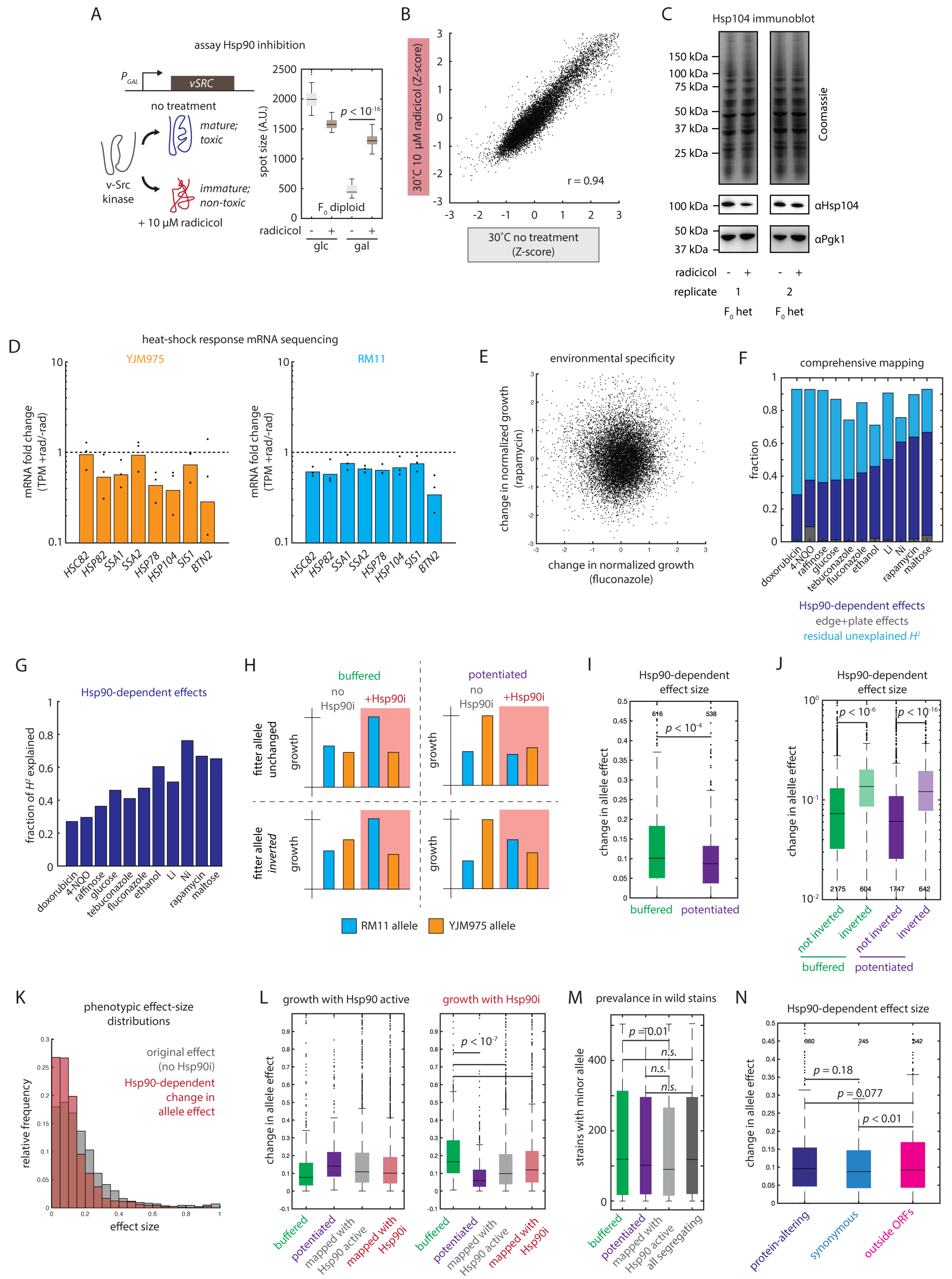
A
assay Hsp90 inhibition
vSRC
no treatment
mature; toxic
immature; non-toxic
v-Src kinase
+ 10 μM radicicol
spot size (A.U.)
F0 diploid
radicicol
glc
gal
B
30˚C 10 μM radicicol (Z-score)
30˚C no treatment (Z-score)
r = 0.94
C
Hsp104 immunoblot
Coomassie
αHsp104
αPgk1
radicicol
replicate
F0 het
D
heat-shock response mRNA sequencing
YJM975
RM11
mRNA fold change (TPM +rad/-rad)
HSC82
HSP82
SSA1
SSA2
HSP78
HSP104
SIS1
BTN2
E
environmental specificity
change in normalized growth (rapamycin)
change in normalized growth (fluconazole)
F
comprehensive mapping
fraction
doxorubicin
4-NQO
raffinose
glucose
tebuconazole
fluconazole
ethanol
Li
Ni
rapamycin
maltose
Hsp90-dependent effects
edge+plate effects
residual unexplained H²
G
Hsp90-dependent effects
fraction of H² explained
H
buffered
potentiated
no Hsp90i
+Hsp90i
fitter allele unchanged
fitter allele inverted
growth
RM11 allele
YJM975 allele
I
Hsp90-dependent effect size
change in allele effect
buffered
potentiated
J
Hsp90-dependent effect size
change in allele effect
not inverted
inverted
buffered
potentiated
K
phenotypic effect-size distributions
original effect (no Hsp90i)
Hsp90-dependent change in allele effect
relative frequency
effect size
L
growth with Hsp90 active
growth with Hsp90i
change in allele effect
mapped with Hsp90 active
mapped with Hsp90i
M
prevalance in wild stains
strains with minor allele
n.s.
all segregating
N
Hsp90-dependent effect size
change in allele effect
protein-altering
synonymous
outside ORFs

3

**Figure S1. To accompany Figure 1.** (A) Growth of hybrid RM11/YJM975 diploid strains bearing pGAL::v-Src in glucose (left) and galactose (right) with Hsp90 active or inhibited by radicicol, as indicated. Data shown are mean and s.e.m.; $n$ = 96 per genotype. (B) Normalized growth of progeny in minimal glucose medium with Hsp90 active (abscissa) and inhibited with radicicol (ordinate). Each dot represents the growth of one segregant. Correlation by Pearsons $r$. (C) Coomassie brilliant blue staining (top), anti-Hsp104 western blot (middle), and anti-Pgk1 western blot (bottom) of hybrid RM11/YJM975 diploid strains propagated with Hsp90 active or inhibited by radicicol, as indicated. $N$ = 3 replicates represent the collection of independent biological replicate cultures. (D) Estimated transcripts per million (TPM) abundance for heat shock-responsive mRNAs in the YJM975 homozygous diploid (left) or RM11 homozygous diploid (right) with Hsp90 active (grey) or inhibited by radicicol (red). Bars show mean; dots show biological replicate cultures; $N$ = 3. (E) Change in normalized growth upon radicicol treatment when progeny were grown in fluconazole (abscissa) and rapamycin (ordinate). Each dot represents the change in normalized growth of one segregant. (F) Fraction of variance in Hsp90-depedent phenotypes attributable to geometric and plate effects (grey), modification of linear genetic effects by Hsp90 activity (dark blue), or unexplained broad-sense heritability (light blue) for growth phenotypes as indicated. (G) Fraction of broad-sense heritability explained by modification of linear genetic effects by Hsp90 for growth phenotypes as indicated. (H) Schematic of buffered and potentiated interactions between a hypothetical variant and Hsp90 in which the sign of the fitness effect of the variant is either preserved or inverted. (I) Absolute change in allele effect for fine-mapped Hsp90-modified variants that were buffered (left; green) or potentiated (right; purple). Box plots show the median and upper and lower quartiles; whiskers show 1.5 times the interquartile range; $p$ value by $t$ test; $n$ as indicated. (J) Absolute change in allele effect for Hsp90-modified variants that were buffered (green, left) or potentiated (purple, right), divided by whether the sign of the fitness effect of the variant was preserved (dark colors) or reversed (light colors). Box plots show the median and upper and lower quartiles; whiskers show 1.5 times the interquartile range; *p* value by *t* test; *n* as indicated. (K) Histograms of effect sizes of linear fine-mapped variants with Hsp90 active (absolute value of normalized growth effect; grey) and fine-mapped Hsp90-modified variants (absolute change in normalized growth effect; red). (L) Absolute effect size for buffered (green) and potentiated (purple) Hsp90-modified variants alongside causal variants mapped with Hsp90 active (grey) and inhibited (red) in environments with Hsp90 active (left) and environments with Hsp90 inhibited (right). *p* values by *t* test. (M) Minor allele frequencies in the 1,002 Yeast Genomes collection for buffered (green) and potentiated (purple) Hsp90-modified variants alongside causal variants mapped with Hsp90 active (light grey) and all segregating variants (dark grey). *p* values by *t* test. (N) Absolute change in allele effect for fine-mapped Hsp90-modified variants that were protein altering (dark blue), synonymous (light blue), and regulatory (pink). Box plots show the median and upper and lower quartiles; whiskers show 1.5 times the interquartile range; *p* values by *t* test; *n* as indicated.

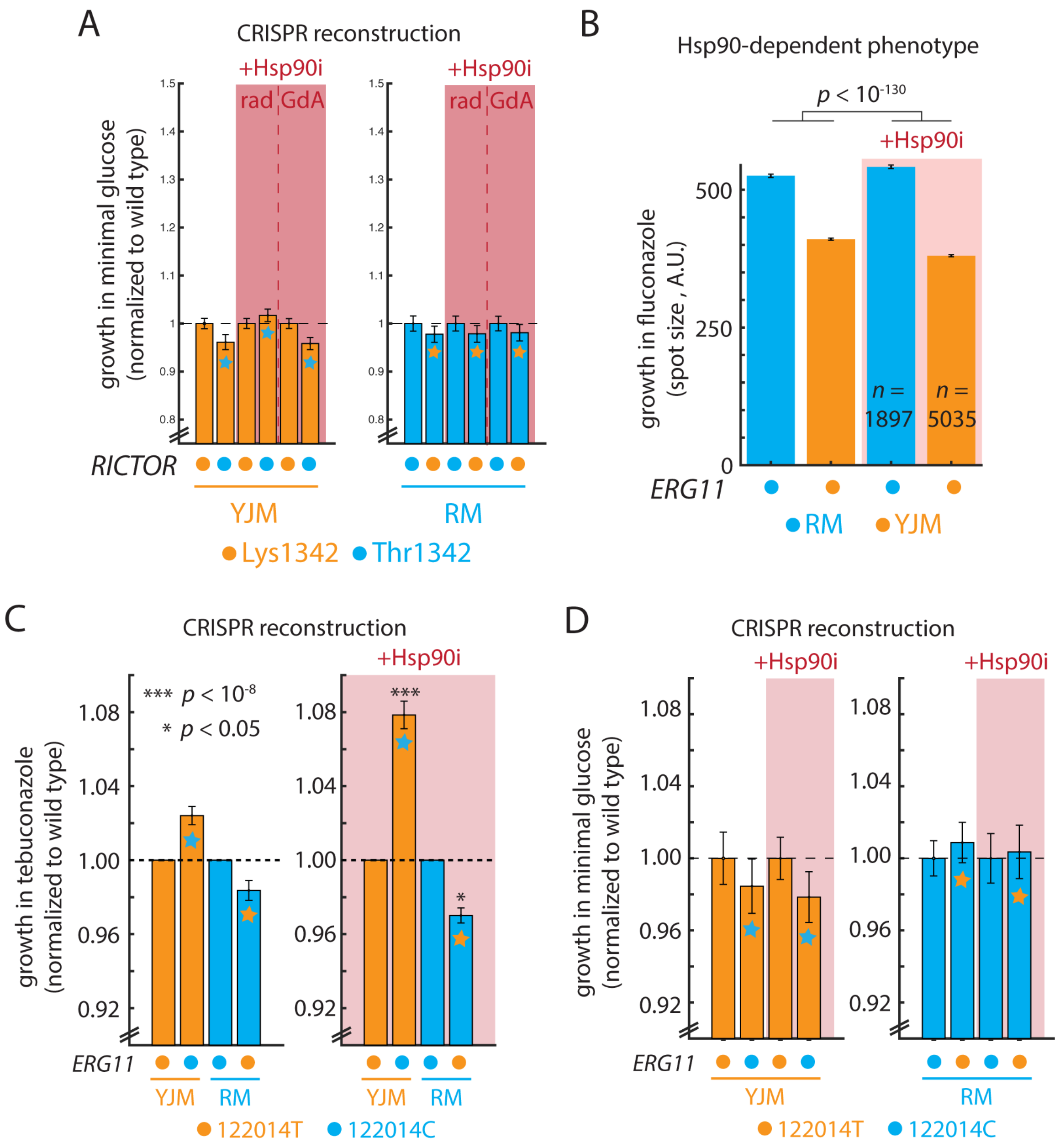


**Figure S2. To accompany Figure 2.** (A) Growth in minimal glucose medium of genome-edited strains with Hsp90 active (no shading) and inhibited (red shading). The left panel shows wild type YJM975 Avo3$^{Lys1342}$ and edited YJM975 Avo3$^{Thr1342}$ strains (blue stars); the right panel shows wild type RM11 Avo3$^{Thr1342}$ strains and edited RM11 Avo3$^{Lys1342}$. Hsp90 was inhibited with radicicol (rad) and geldanamycin A (GdA). Data shown are mean and s.e.m. normalized to the wild type; *n* = 96 per genotype. (B) Growth in fluconazole with (red shading) and without (no shading) radicicol of progeny bearing the resistant (blue) and sensitive (orange) alleles of *ERG11*. Data shown are mean and s.e.m. for the indicated number of progeny with each homozygous genotype; *F* test *p*-value from multivariate regression. (C) Growth in tebuconazole of genome-edited strains with Hsp90 active (no shading) and inhibited with radicicol (+Hsp90i; red shading). Each panel shows wild type YJM975 *ERG11*$^{122014T}$, edited YJM975 *ERG11*$^{122014C}$ (blue stars), wild type RM11 *ERG11*$^{122014C}$, and edited RM11 *ERG11*$^{122014T}$ (orange stars) strains. Data shown are mean and s.e.m. normalized to the wild type; *p* values by *t* test against the same mutant strain without radicicol treatment; *n* = 384 per genotype. (D) Growth in minimal glucose medium of genome-edited strains with Hsp90 active (no shading) and inhibited with radicicol (+Hsp90i; red shading). Each panel shows wild type YJM975 *ERG11*$^{122014T}$, edited YJM975 *ERG11*$^{122014C}$ (blue stars), wild type RM11 *ERG11*$^{122014C}$, and edited RM11 *ERG11*$^{122014T}$ (orange stars) strains. Data shown are mean and s.e.m. normalized to the wild type; *n* = 384 per genotype.

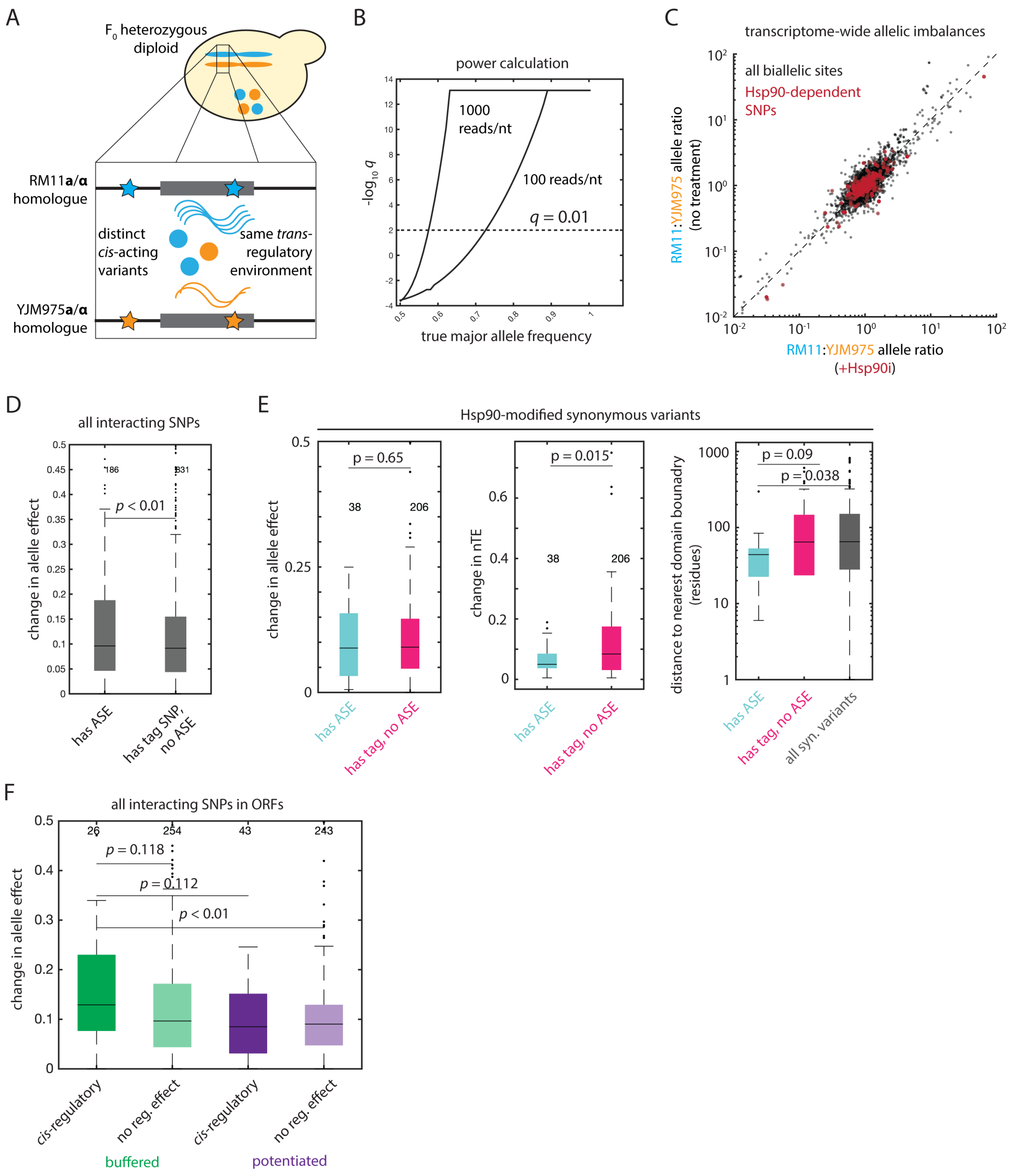


**Extended Data Figure 3. To accompany Figure 3.** (A) Schematic of allele-specific expression analysis in the hybrid RM11/YJM975 diploid. (B) Estimated Bonferroni-corrected $q$ value (ordinate) for allele-specific expression as a function of true major allele frequency (abscissa) for read depths of 100 and 1000 counts per nucleotide. (C) Allelic ratio (RM counts/YJM975 counts) in the hybrid diploid with Hsp90 inhibited by radicicol (abscissa) or with Hsp90 active (ordinate) for all biallelic SNPs (grey) and for SNPs with Hsp90-dependent phenotypic effects (red). Data shown are mean allelic ratios across three replicates for each condition. (D) Absolute change in allele effect for all Hsp90-modified variants that exhibited allele-specific expression (left) or had a tag SNP but no allele-specific expression (right). Box plots show the median and upper and lower quartiles; whiskers show 1.5 times the interquartile range; *p* value by *t* test; *n* as indicated. (E) Absolute change in allele effect (left), absolute change in normalized translation efficiency (nTE), and distance to the nearest protein domain boundary from SuperFam (right) for Hsp90-dependent synonymous variants with allele-specific expression (blue), no ASE (pink), and all segregating synonymous variants (grey), as indicated. (F) Absolute change in allele effect for Hsp90-modified variants within ORFs that were buffered (green) and potentiated (purple), sorted by whether the transcript exhibited allele-specific expression (left) or had a tag SNP but no allele-specific expression (right). Box plots show the median and upper and lower quartiles; whiskers show 1.5 times the interquartile range; *p* values by *t* test; *n* as indicated.

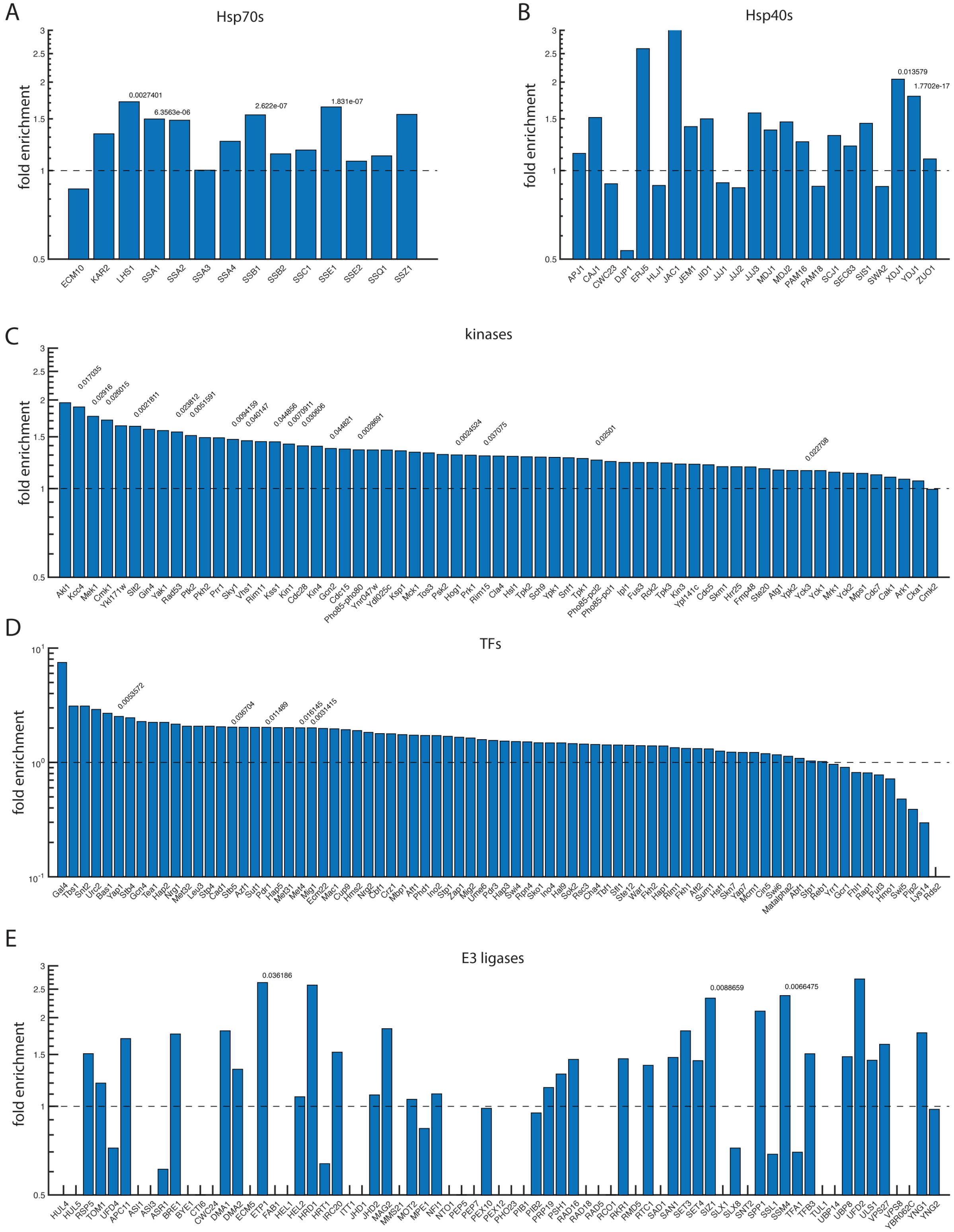
A
Hsp70s
fold enrichment
0.0027401
6.3563e-06
2.622e-07
1.831e-07
ECM10 KAR2 LHS1 SSA1 SSA2 SSA3 SSA4 SSB1 SSB2 SSC1 SSE1 SSE2 SSQ1 SSZ1
B
Hsp40s
fold enrichment
0.013579
1.7702e-17
APJ1 CAJ1 CWC23 DJP1 ERJ5 HLJ1 JAC1 JEM1 JID1 JJJ1 JJJ2 JJJ3 MDJ1 MDJ2 PAM16 PAM18 SCJ1 SEC63 SIS1 SWA2 XDJ1 YDJ1 ZUO1
C
kinases
fold enrichment
Akl1 Kcc4 Mek1 Cmk1 Ykl171w Stt2 Gin4 Yak1 Rad53 Ptk2 Pkh2 Prr1 Sky1 Vhs1 Rim11 Kss1 Kin1 Cdc28 Kin4 Gcn2 Cdc15 Pho85-pho80 Ynr047w Ydl025c Ksp1 Mck1 Tos3 Psk2 Hog1 Prk1 Rim15 Cla4 Hsl1 Tpk2 Sch9 Ypk1 Snf1 Tpk1 Pho85-pcl2 Pho85-pcl1 Ipl1 Fus3 Rck2 Tpk3 Kin3 Ypl141c Cdc5 Skm1 Hrr25 Fmp48 Ste20 Atg1 Ypk2 Yck3 Yck1 Mrk1 Yck2 Mps1 Cdc7 Cak1 Ark1 Cka1 Cmk2
D
TFs
fold enrichment
E
E3 ligases
fold enrichment
0.036186
0.0088659
0.0066475
HUL4 HUL5 RSP5 TOM1 UFD4 APC11 ASI1 ASI3 ASR1 BRE1 BYE1 CTI6 CWC24 DMA1 DMA2 ECM5 ETP1 FAB1 HEL1 HEL2 HRD1 HRT1 IRC20 ITT1 JHD1 JHD2 MAG2 MMS21 MOT2 MPE1 NFI1 NTO1 PEP5 PEP7 PEX10 PEX12 PHO23 PIB1 PIB2 PRP19 PSH1 RAD16 RAD18 RAD5 RCO1 RKR1 RMD5 RTC1 SAD1 SAN1 SET3 SET4 SIZ1 SLX1 SLX8 SNT2 SPP1 SSL1 SSM4 TFA1 TFB3 TUL1 UBP14 UBR1 UFD2 ULS1 VPS27 VPS8 YBR062C YNG1 YNG2

81

**Figure S4. To accompany Figure 4.** Relative enrichment of interactors or targets of the indicated proteins amongst Hsp90-modified alleles as compared to all segregating genetic variants for (A) Hsp70 chaperones; (B) Hsp40 chaperones; (C) all transcription factors; (D) all kinases; and (E) all E3 ligases. Bonferroni-corrected *q* values from Fisher's exact test.

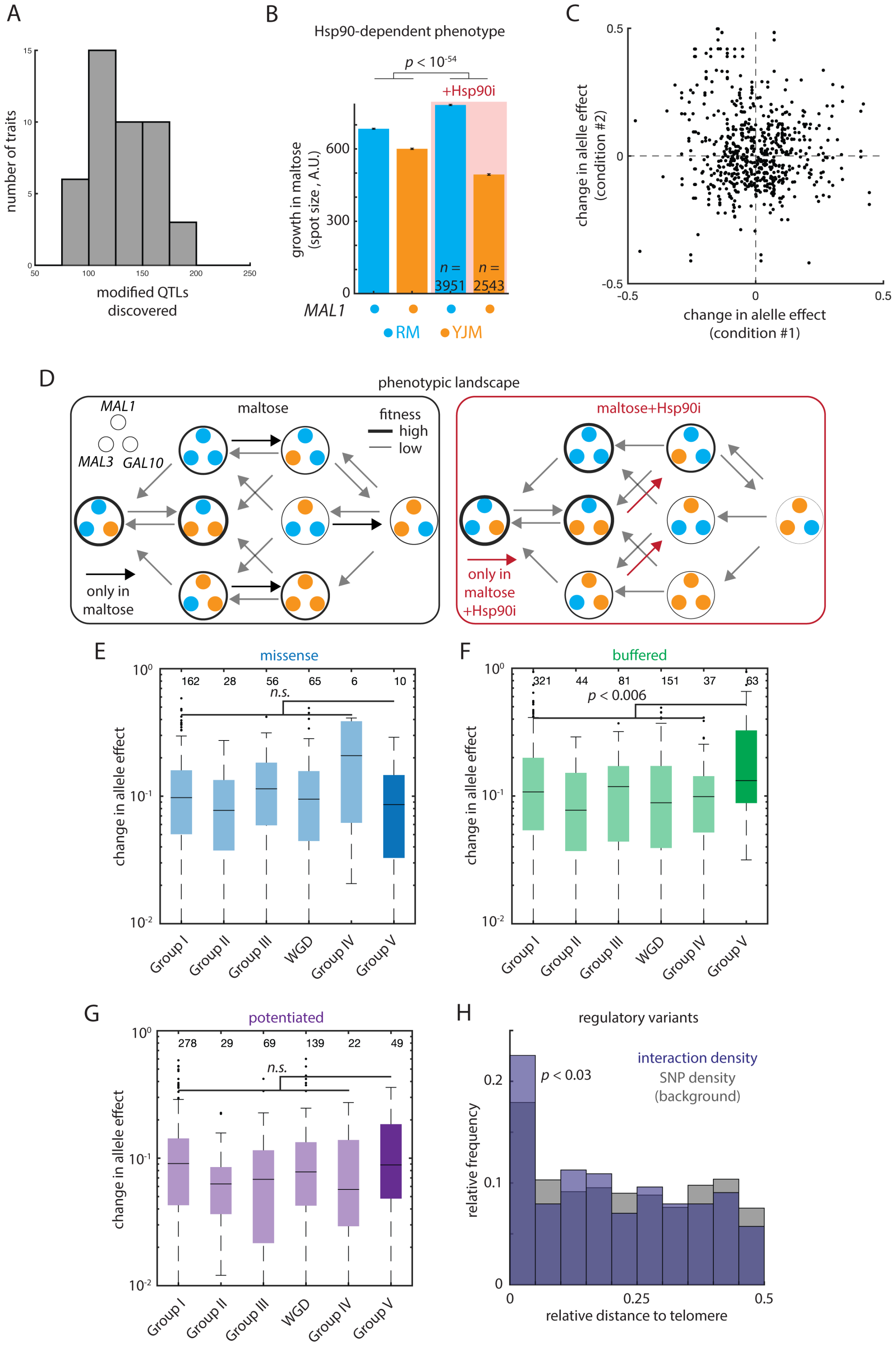
A
number of traits
modified QTLs discovered
B
Hsp90-dependent phenotype
p < 10-54
+Hsp90i
growth in maltose (spot size , A.U.)
n = 3951
n = 2543
MAL1
RM
YJM
C
change in alelle effect (condition #2)
change in alelle effect (condition #1)
D
phenotypic landscape
MAL1
MAL3
GAL10
maltose
fitness
high
low
only in maltose
maltose+Hsp90i
only in maltose +Hsp90i
E
missense
change in allele effect
n.s.
Group I
Group II
Group III
WGD
Group IV
Group V
F
buffered
p < 0.006
G
potentiated
n.s.
H
regulatory variants
p < 0.03
interaction density
SNP density (background)
relative frequency
relative distance to telomere

86

**Figure S5. To accompany Figure 5 and Figure 6.** (A) Histogram of the number of Hsp90-dependent loci discovered for each phenotype. (B) Normalized growth in maltose with Hsp90 active (no shading) and inhibited with radicicol (red shading) of progeny with genotypes as indicated (RM11, blue; YJM975, orange) at *MAL1*. Data shown are mean and s.e.m. (C) Scatterplot of the change in allele effect upon Hsp90 inhibition in one environment (abscissa) compared to the change in allele effect for the same modified variant identified in a second environment (ordinate). Shown are all possible environment-environment pairs for modified variants mapped in multiple conditions. (D) Phenotypic landscape of growth in maltose (left) or maltose with radicicol (right). Genotypes at *MAL1, MAL3,* and *GAL10* are as indicated; the fitness of each genotype is encoded in the line width of the enclosing circle. Gray arrows indicate permissible mutations that do not significantly decrease fitness (nominal $p > 0.05$) in either condition; black arrows show mutations only permissible in maltose; red arrows show mutations only permissible in maltose with radicicol. Absolute change in allele effect for (E) protein-coding Hsp90-modified variants; (F) buffered Hsp90-modified variants; and (G) potentiated Hsp90-modified variants with ages as indicated. Box plots show the median and upper and lower quartiles; whiskers show 1.5 times the interquartile range; $p$ value by $t$ test of the indicated age group against all other variants. $n$ as indicated. (H) Relative frequency of Hsp90-modified regulatory variants (blue) and all segregating regulatory variants (grey) as a function of distance from the nearest chromosome end (normalized to chromosome length). $p$ value by Kolmogorov-Smirnov test.

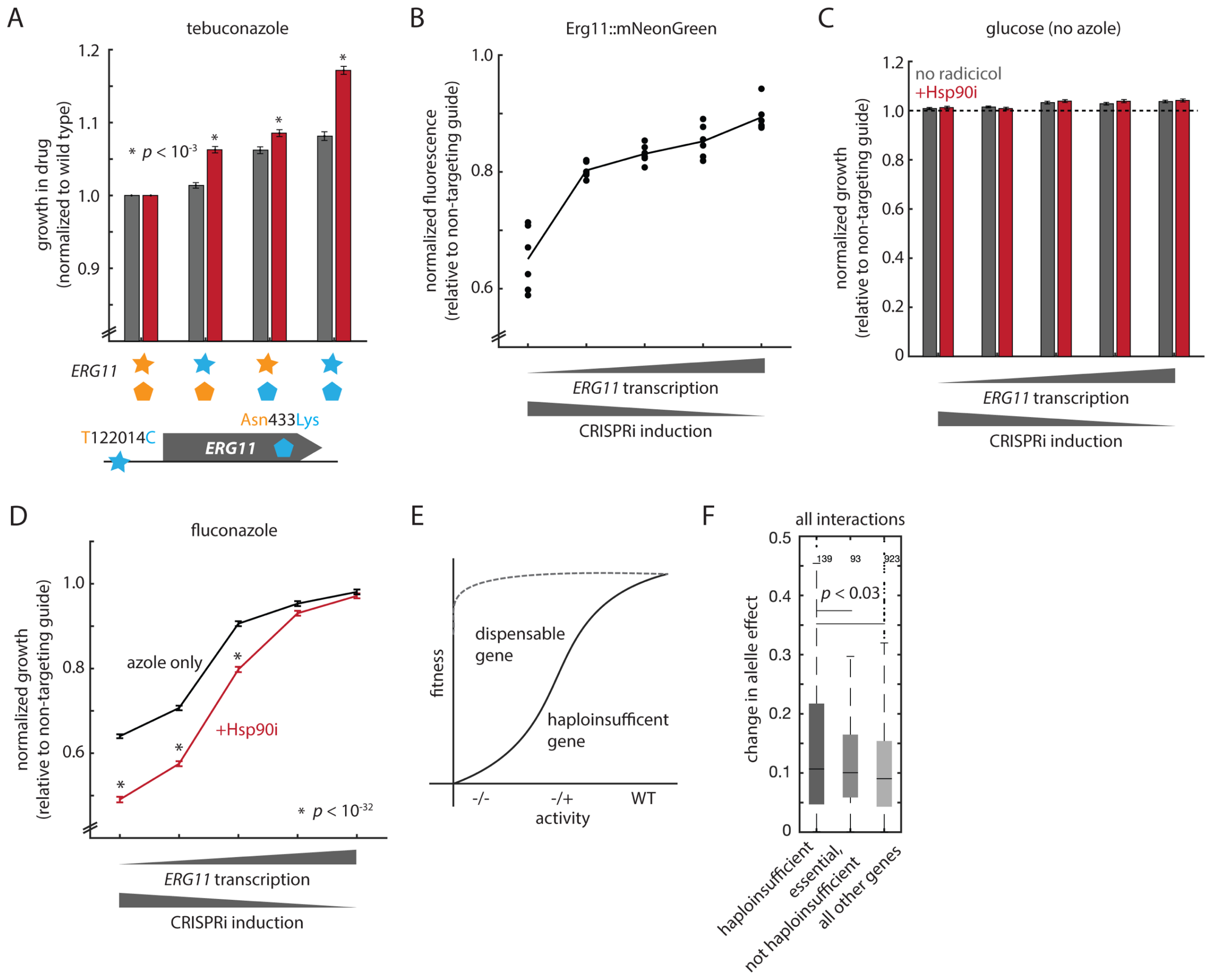


**Figure S6. To accompany Figure 7.** (A) Growth in tebuconazole of genome-edited strains with Hsp90 active (grey) and inhibited with radicicol (+Hsp90i; red). Each panel shows wild type YJM975, edited YJM975 *ERG11*$^{122014C}$, edited YJM975 Erg11$^{433Lys}$, and edited double-mutant YJM975 *ERG11*$^{122014C}$ Erg11$^{433Lys}$ strains. Data shown are mean and s.e.m. normalized to the wild type; *p* values by *t* test against the same mutant strain without radicicol treatment; $n = 384$ per genotype. (B) Erg11::mNeonGreen fusion fluorescence of BY4741 haploid strains expressing an anti-*ERG11* gRNA normalized to strains expressing a non-targeting control gRNA. Shown is mean of $n = 6$ replicates; dots show each replicate. The concentrations of inducing molecule (aTc) added to each culture are (left to right) 1000 ng/μL, 500 ng/μL, 250 ng/μL, 125 ng/μL, and 62.5 ng/μL

(C) Growth in minimal glucose of YJM975 haploid strains expressing an anti-*ERG11* gRNA normalized to strains expressing a non-targeting control gRNA without the addition of radicicol (grey) or with radicicol (+Hsp90i; red). The concentrations of inducing molecule (aTc) added to each culture are (left to right) 1000 ng/µL, 500 ng/µL, 250 ng/µL, 125 ng/µL, and 62.5 ng/µL. Data shown are mean and s.e.m. normalized to the non-targeting guide; $n$ = 384 per genotype. (D) Growth of RM11 haploid strains expressing an anti-*ERG11* gRNA (closed boxes) normalized to the growth of the YJM975 strain expressing the non-targeting control gRNA. Growth is in fluconazole without the addition of radicicol (black) or with radicicol (+Hsp90i; red). The concentrations of inducing molecule (aTc) added to each culture are (left to right) 1000 ng/µL, 500 ng/µL, 250 ng/µL, 125 ng/µL, and 62.5 ng/µL. (E) Schematic of the activity-fitness relationship for a dispensable and a haploinsufficient gene. (F) Absolute change in allele effect for Hsp90-modified variants in haploinsufficient genes, essential but not haploinsufficient genes, or all other genes, as indicated. Box plots show the median and upper and lower quartiles; whiskers show 1.5 times the interquartile range; $p$ values by $t$ test; $n$ as indicated.

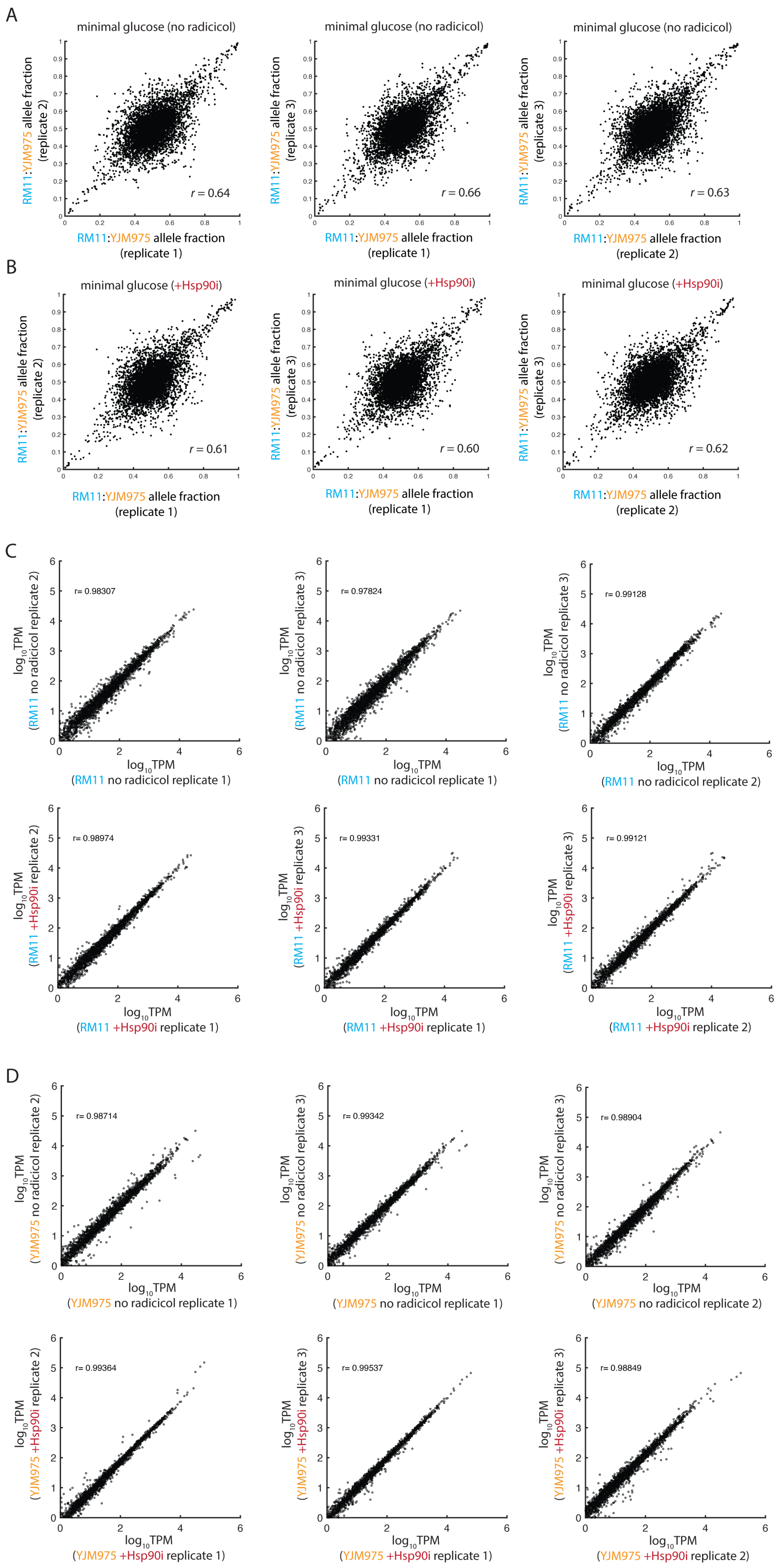
A
minimal glucose (no radicicol)
RM11:YJM975 allele fraction (replicate 2)
RM11:YJM975 allele fraction (replicate 1)
r = 0.64
minimal glucose (no radicicol)
RM11:YJM975 allele fraction (replicate 3)
RM11:YJM975 allele fraction (replicate 1)
r = 0.66
minimal glucose (no radicicol)
RM11:YJM975 allele fraction (replicate 3)
RM11:YJM975 allele fraction (replicate 2)
r = 0.63
B
minimal glucose (+Hsp90i)
RM11:YJM975 allele fraction (replicate 2)
RM11:YJM975 allele fraction (replicate 1)
r = 0.61
minimal glucose (+Hsp90i)
RM11:YJM975 allele fraction (replicate 3)
RM11:YJM975 allele fraction (replicate 1)
r = 0.60
minimal glucose (+Hsp90i)
RM11:YJM975 allele fraction (replicate 3)
RM11:YJM975 allele fraction (replicate 2)
r = 0.62
C
log10TPM (RM11 no radicicol replicate 2)
log10TPM (RM11 no radicicol replicate 1)
r= 0.98307
log10TPM (RM11 no radicicol replicate 3)
log10TPM (RM11 no radicicol replicate 1)
r= 0.97824
log10TPM (RM11 no radicicol replicate 3)
log10TPM (RM11 no radicicol replicate 2)
r= 0.99128
log10TPM (RM11 +Hsp90i replicate 2)
log10TPM (RM11 +Hsp90i replicate 1)
r= 0.98974
log10TPM (RM11 +Hsp90i replicate 3)
log10TPM (RM11 +Hsp90i replicate 1)
r= 0.99331
log10TPM (RM11 +Hsp90i replicate 3)
log10TPM (RM11 +Hsp90i replicate 2)
r= 0.99121
D
log10TPM (YJM975 no radicicol replicate 2)
log10TPM (YJM975 no radicicol replicate 1)
r= 0.98714
log10TPM (YJM975 no radicicol replicate 3)
log10TPM (YJM975 no radicicol replicate 1)
r= 0.99342
log10TPM (YJM975 no radicicol replicate 3)
log10TPM (YJM975 no radicicol replicate 2)
r= 0.98904
log10TPM (YJM975 +Hsp90i replicate 2)
log10TPM (YJM975 +Hsp90i replicate 1)
r= 0.99364
log10TPM (YJM975 +Hsp90i replicate 3)
log10TPM (YJM975 +Hsp90i replicate 1)
r= 0.99537
log10TPM (YJM975 +Hsp90i replicate 3)
log10TPM (YJM975 +Hsp90i replicate 2)
r= 0.98849

**Figure S7. To accompany Method Details.** Cross-plotted are RM11:YJM975 allelic ratios for all biallelic SNPs from three biological replicate cultures of the F0 heterozygous RM11/YJM975 diploid grown without Hsp90 inhibition (A) or with radicicol (+Hsp90i; B). (C) Cross-plotted are mRNA TPM estimates from three biological replicate RM11 homozygous diploid cultures grown without Hsp90 inhibition (top) or with radicicol (+Hsp90i; bottom). (D) As in (A), but for YJM975 homozygous diploids.

**Supplemental Table S1:** Stress conditions used in this study.

| **Compound** | **Final concentration** |
|---|---|
| maltose | 2% [no glucose added] |
| raffinose | 2% [no glucose added] |
| ethanol | 2% [no glucose added] |
| 4-nitroquinoline 1-oxide [4-NQO] | 4 μM |
| doxorubicin | 40 μM |
| fluconazole | 100 μM |
| lithium [as chloride; $Li^{+}$] | 10 mM |
| nickel [as sulfate; $Ni^{2+}$] | 1 mM |
| rapamycin | 5 μM |
| tebuconazole | 0.6 μM |

The standard media formulation was 2% glucose [Fisher], 6.7 g/L yeast nitrogen base [RPI], and 20 mg/L uracil [Sigma], with 2% agar [IBI] included for solid media. Anhydrotetracycline [Abcam] was formulated at 5 g/L in 85% ethanol.
